# Whole genome sequencing for diagnosis of neurological repeat expansion disorders

**DOI:** 10.1101/2020.11.06.371716

**Authors:** Kristina Ibanez, James Polke, Tanner Hagelstrom, Egor Dolzhenko, Dorota Pasko, Ellen Thomas, Louise Daugherty, Dalia Kasperaviciute, Ellen M McDonagh, Katherine R Smith, Antonio Rueda Martin, Dimitris Polychronopoulos, Heather Angus-Leppan, Kailash P Bhatia, James E Davison, Richard Festenstein, Pietro Fratta, Paola Giunti, Robin Howard, Laxmi Venkata Prasad Korlipara, Matilde Laurá, Meriel McEntagart, Lara Menzies, Huw Morris, Mary M Reilly, Robert Robinson, Elisabeth Rosser, Francesca Faravelli, Anette Schrag, Jonathan M Schott, Thomas T Warner, Nicholas W Wood, David Bourn, Kelly Eggleton, Robyn Labrum, Philip Twiss, Stephen Abbs, Liana Santos, Ghareesa Almheiri, Isabella Sheikh, Jana Vandrovcova, Christine Patch, Ana Lisa Taylor Tavares, Zerin Hyder, Anna Need, Helen Brittain, Emma Baple, Loukas Moutsianas, Genomics England Research Consortium, Viraj Deshpande, Denise L Perry, Shankar Ajay, Aditi Chawla, Vani Rajan, Kathryn Oprych, Patrick F Chinnery, Angela Douglas, Gill Wilson, Sian Ellard, Karen Temple, Andrew Mumford, Dom McMullan, Kikkeri Naresh, Frances Flinter, Jenny C Taylor, Lynn Greenhalgh, William Newman, Paul Brennan, John A. Sayer, F Lucy Raymond, Lyn S Chitty, Zandra C Deans, Sue Hill, Tom Fowler, Richard Scott, Henry Houlden, Augusto Rendon, Mark J Caulfield, Michael A Eberle, Ryan J Taft, Arianna Tucci

## Abstract

**Background:** Repeat expansion (RE) disorders affect ~1 in 3000 individuals and are clinically heterogeneous diseases caused by expansions of short tandem DNA repeats. Genetic testing is often locus-specific, resulting in under diagnosis of atypical clinical presentations, especially in paediatric patients without a prior positive family history. Whole genome sequencing (WGS) is emerging as a first-line test for rare genetic disorders, but until recently REs were thought to be undetectable by this approach.

**Methods:** WGS pipelines for RE disorder detection were deployed by the 100,000 Genomes Project and Illumina Clinical Services Laboratory. Performance was retrospectively assessed across the 13 most common neurological RE loci using 793 samples with prior orthogonal testing (182 with expanded alleles and 611 with alleles within normal size) and prospectively interrogated in 13,331 patients with suspected genetic neurological disorders.

**Findings:** WGS RE detection showed minimum 97·3% sensitivity and 99·6% specificity across all 13 disease-associated loci. Applying the pipeline to patients from the 100,000 Genomes Project identified pathogenic repeat expansions which were confirmed in 69 patients, including seven paediatric patients with no reported family history of RE disorders, with a 0.09% false positive rate.

**Interpretation:** We show here for the first time that WGS enables the detection of causative repeat expansions with high sensitivity and specificity, and that it can be used to resolve previously undiagnosed neurological disorders. This includes children with no prior suspicion of a RE disorder. These findings are leading to diagnostic implementation of this analytical pipeline in the NHS Genomic Medicine Centres in England.

**Funding:** Medical Research Council, Department of Health and Social Care, National Health Service England, National Institute for Health Research, Illumina Inc

## INTRODUCTION

Despite recent advances in our understanding of the genetic basis of rare neurological disorders, ~70% of patients remain genetically undiagnosed.^1^ This is partly attributable to undertesting of genetic variants such as repeat expansions (RE), which are a leading cause of over 40 neurological disorders.^2^ RE disorders include the most common neurogenetic conditions, such as Huntington disease (HD), amyotrophic lateral sclerosis (ALS), frontotemporal dementia (FTD), and Fragile X syndrome, a common cause of intellectual disability.^2^ RE disorders are clinically and genetically heterogeneous. The same repeat expansion can be associated with different phenotypes, within the same family. For example, *C9orf72* is associated with both amyotrophic lateral sclerosis (ALS) and frontotemporal dementia (FTD).^3^ Furthermore, REs in different loci can present with overlapping phenotypic features, such as the spinocerebellar ataxia (SCA) genes, can present as an autosomal dominant cerebellar ataxia.^4^

RE disorders are associated with an increase in the number of repetitive short tandem DNA sequences, and the pathogenicity thresholds for each disorder are locus specific. These repeats exhibit molecular instability which can lead to changes in size across generations (generally increasing in length) and tissues.^5^ In these conditions, increases in the number of repeats often lead to earlier onset and more severe disease in successive generations within the same family.^2^ Paediatric onset of RE disorders can present as multi-system syndromes without specific phenotypic signatures,^6^ and are therefore more likely to be under-diagnosed and under-tested due phenotypic overlap with other early-onset genetic disorders.^7^

Laboratory assessment of REs includes targeted molecular assessment of individual loci, guided by the clinical diagnosis, using PCR-based or Southern blot^8^ assays which can be costly and time-consuming. Additionally, due to the the varied and overlapping phenotypic features of these disorders, most RE loci remain untested in an undiagnosed individual.^9^

Whole genome sequencing (WGS) is emerging as a first-line diagnostic tool in defined cases of rare disease,^10^ but was previously thought to have limited capability to assess highly repetitive repeat expansion loci.^11^ Here, we report on the validation and deployment of a RE-aware WGS pipeline as part of the 100,000 Genomes Project (GE) and the Illumina Clinical Services Laboratory (ICSL), and its application to patients with undiagnosed neurological disorders (**Figure 1**).

**Figure 1.**
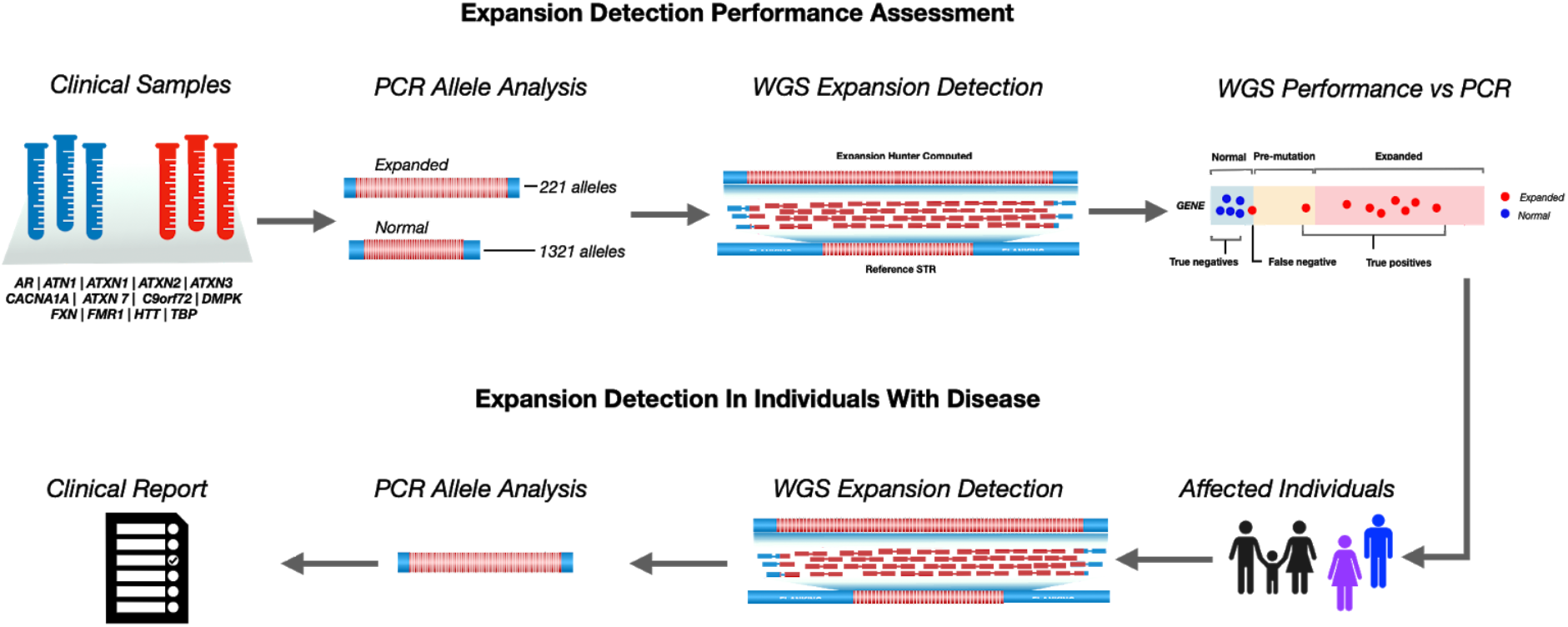
Evaluation of WGS repeat expansion detection performance and its application to patients with genetically undiagnosed disorders. In this study, thirteen well-established repeat expansion disorders were selected for interrogation by whole genome sequencing. Performance was assessed against 221 expanded and 1321 normal alleles drawn from samples tested at Neurogenetics Laboratory at the National Hospital for Neurology and Neurosurgery and Genetics Laboratory, Cambridge University Hospitals NHS Foundation Trust (see **Methods and Supplemental Methods**). WGS was performed at a minimum of 35x depth and each disease locus was interrogated using the Expansion Hunter software package (see **Methods and Supplemental Methods**). Expansion detection performance for each locus was assessed against the pre-mutation cutoff (**Table S3**). Within this dataset we observed an overall sensitivity of 98·2% and specificity of 99·9%, which when applied to individuals with a genetically undiagnosed disorder from the 100,000 Genomes Project (Genomics England, GE), or tested by the Illumina Clinical Services Laboratory (ICSL) revealed previously undetected expansions in *AR*, *ATN1*, *ATXN1*, *ATXN2*, *ATXN3*, *ATXN7*, *CACNA1A*, *C9orf72*, *HTT*, *TBP*, *DMPK,* and *FXN*.

## METHODS

### Whole genome sequencing

DNA was prepared for sequencing using TruSeq DNA PCR-Free library preparation and 150 or 125 bp paired-end sequencing was performed on either HiSeq 2000 or HiSeq X platforms. Genomes were sequenced to an average minimum depth of 35X (31X - 37X) (**Table S1**).

### Repeat expansion performance datasets

WGS RE performance was evaluated using data from two sources: 254 participants from the 100,000 Genomes Project at Genomics England (GE), and 150 individuals previously tested for expansions as part of clinical assessment from the NHS Genomic Laboratory based at Cambridge University Hospitals NHS Foundation Trust and used for External UK National Quality Assurance Schemes and sequenced in ICSL (**Table S2**, and **Supplementary methods**).

### PCR analysis

RE were assessed by polymerase chain reaction (PCR) amplification and fragment analysis; Southern blotting was performed for large *C9orf72* expansions. For additional details, including primer sequences see **Supplementary methods**.

### Repeat expansion genotyping and visual inspection

Short tandem repeat (STR) genotyping from whole genome sequencing was performed using the ExpansionHunter (EH) software package.^12,13^ In brief, EH assembles sequencing reads across a pre-defined set of STRs using both mapped and unmapped reads (with the repetitive sequence of interest) to estimate the size of both alleles from an individual (see **Supplementary Methods)**. Recent guidelines from the Association for Medical Pathology and the College of American Pathologists recommend visual inspection of variant calls during routine sign out of NGS variants.^14^ However, short tandem repeat variants cannot be adequately visualised by common visualization tools such as IGV.^15^ To examine WGS data underlying each genotype call, we used a tool that creates a static visualization of the WGS reads containing the repeat identified by EH and used to support the repeat size estimate at each allele. This graph enables direct visualization of haplotypes and the corresponding read pileup of the EH genotypes (https://github.com/Illumina/GraphAlignmentViewer; **Figure 2C and 2D**). Visual inspection of the pileup graph was performed on all WGS-STR calls to (i) confirm the EH prediction for alleles entirely contained in each read (i.e. smaller than the sequencing read length); (ii) confirm the presence of a monoallelic or biallelic expansion; (iii) detect putative false positive calls; (iii) detect false negative alleles in biallelic repeat expansions, such as *FXN* (**Supplementary methods, Figure S1**).

**Figure 2.**
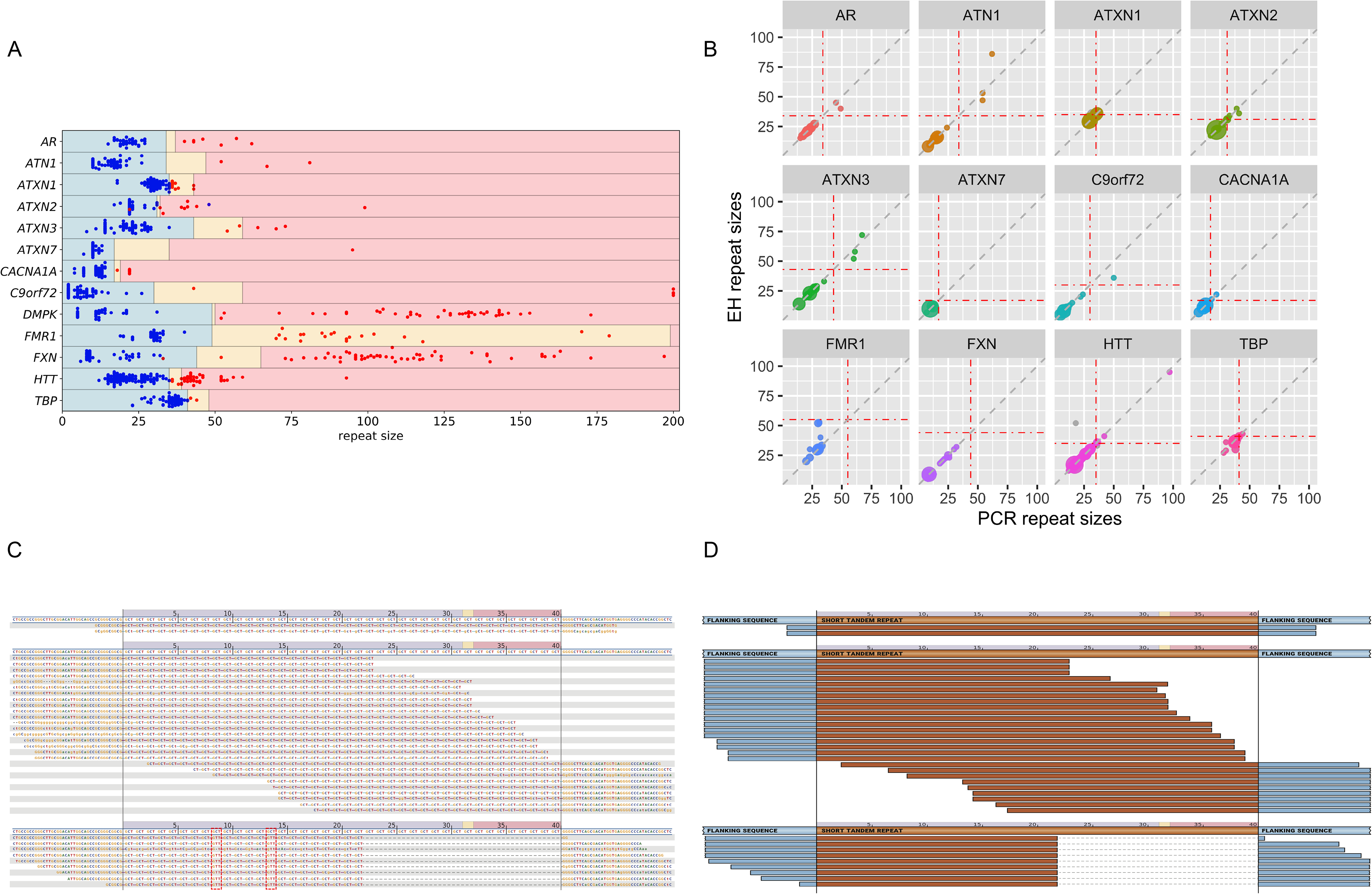
Repeat expansion detection performance using whole genome sequencing. **A) Swim lane plot.** Sizes of repeat-unit expansions predicted by ExpansionHunter across 793 expansion calls. Each genome assessed is represented by two points corresponding to each allele for each locus, with the exception of those on chromosome X (i.e. *FMR1*, *AR*) in males. Points indicate the sequencing based repeat length and the colors indicate the repeat size as assessed by PCR (blue = PCR-normal; red = PCR-expanded). The regions are shaded to indicate normal (blue), premutation (yellow), and expansion (light red) ranges for each gene as indicated in **Table S3**. Blue points in yellow or red shaded regions indicate false positives and the red points in blue shaded regions indicate false negatives. There are five genomes with ~500 repeats in *C9orf72* that are shown here as 200 to facilitate reasonable X axis scaling. The individual calls are provided in **Table S4**. **B) Repeat-size accuracy split by locus**. Bubble-plot representing PCR and EH repeat-sizes in X and Y axis respectively, and the size of each dot showing the number of cases with the same repeat-size. There are two layers, one in grey, and the other colored. The difference between them is the values of the Y axis, being the EH estimations before visual inspection for the grey scenario, and the corrected EH sizes after visual inspection for the coloured layer. Vertical and horizontal red dot dashed lines represent the premutation cut-off (**Table S3**) for each locus. **C) Characteristic pileup graph.** A characteristic pileup graph illustrating a call in *ATXN2* where the estimated genotype for ‘GCT’ repeat unit is 22/40. Reads supporting each genotype are grouped based on the predicted genotype, in this example in three groups: i) two reads supporting 40 repeat units, in the pathogenic range, on the top of the graph; ii) reads flanking the repeat, supporting > 39 repeat units, in the middle; iii) nine reads supporting 22 repeat units, bottom of the graph. **D) Schematic representation of the pileup graph.** Each read has been coloured according to its sequence content, with blue representing the sequence flanking the repeat, and brown the repeated sequence.

### 100,000 Genomes Project patient inclusion criteria

The 100,000 Genomes Project is a UK programme combining diagnostic discovery through research and clinical implementation to assess the value of WGS in individuals with unmet diagnostic needs in rare disease and cancer (**Supplementary Methods**). Following ethical approval (14/EE/1112) participants were enrolled in the 100,000 Genomes Project if they or their guardian had given research consent (n=91,290). They were recruited by healthcare professionals and researchers as part of the Rare Disease cohort (n=35,653) drawn from 13 Genomic Medicine Centres funded and established by NHS England in the National Health Service (NHS) in England. In this study participants with neurological phenotypes (n=13,331) were assessed for RE expansions.

## RESULTS

### Performance of the pipeline

Thirteen pathogenic repeats, that represent a broad spectrum of the most common neurological repeat expansion disorders, were selected for performance assessment (**Table S2**). Specifically, eleven repeat expansion loci associated with ataxia and late-onset neurodegenerative disorders (all ‘CAG’ repeats: *HTT, AR, ATN1, ATXN1, ATXN2, ATXN3, ATXN7, CACNA1A,* and *TBP* plus *C9orf72* and *FXN*), one locus associated with intellectual disability (*FMR1*) and one locus associated with myotonic dystrophy (*DMPK*). Each of these thirteen repeats had PCR validation data for at least one expanded allele (**Table S2**).

#### Detection of expanded alleles

In practice, repeat expansion detection requires accurate categorization of alleles into normal and expanded size ranges. To assess this, a combined set of 793 patient PCR tests, comprising 1,321 normal alleles (below premutation as detailed in **Table S3**) and 221 expanded alleles, covering all 13 disease loci, was established by GE and ICSL (**Table S2**). Comparing the EH output against this benchmark dataset showed a correct classification of 215 out of 221 expanded alleles and of 1,316 out of 1,321 normal alleles (**Table S4, Table S5**), showing a total sensitivity of 97·3% (95% CI: 94·2%-99%) and specificity of 99·6% (95% CI:99·1% - 99·9%) (**Table 1**). All calls were visually inspected and re-classified as appropriate based on the quality of the reads supporting each call (**Supplementary methods**). Following the visual correction, sensitivity was 99·1% (95% CI: 96·7%-99·9%) and specificity of 100% (95% CI:99·7% - 100%) (**Figure 2**; **Table 1**). We note that visualization of the expanded calls was able to detect false positives and to re-classify all false negative alleles in *FXN*, where only one allele was classified as expanded in samples with biallelic expansions (see **Supplementary methods**, **Figure S2**, **Figure S3)**.

**Table 1.**
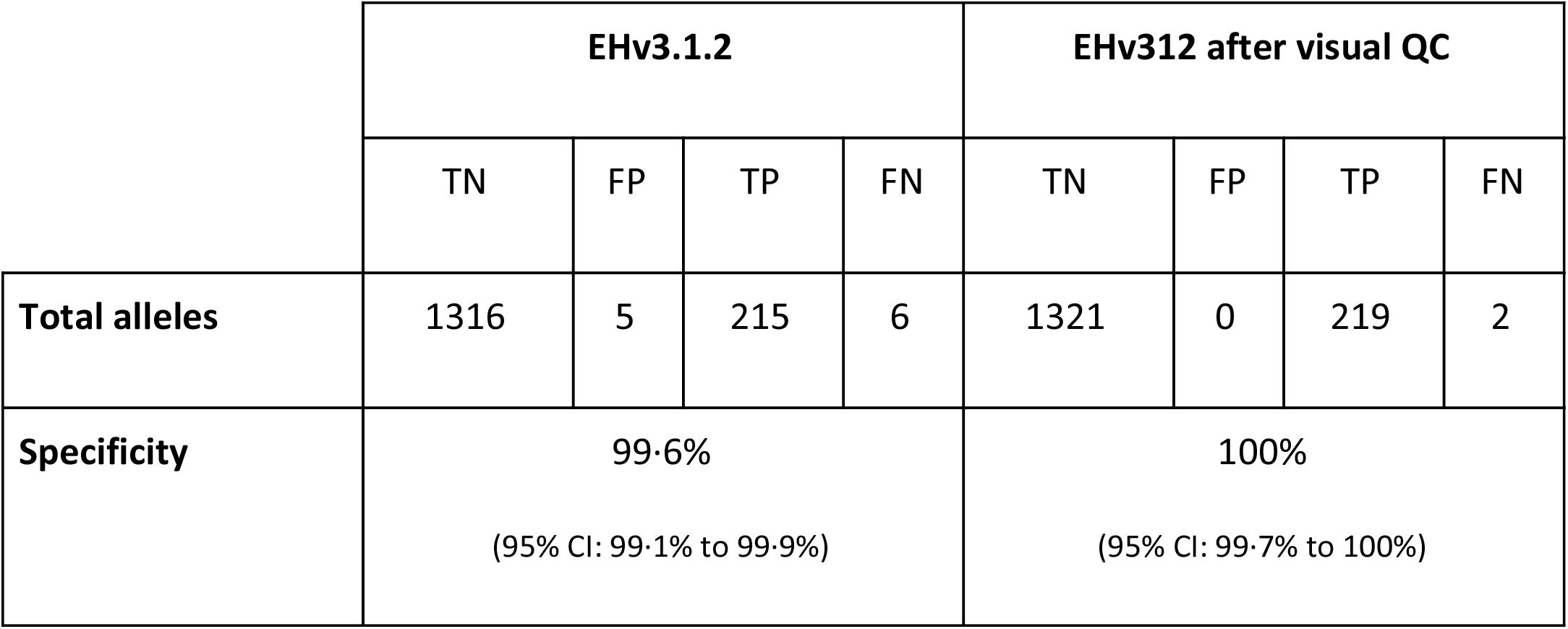
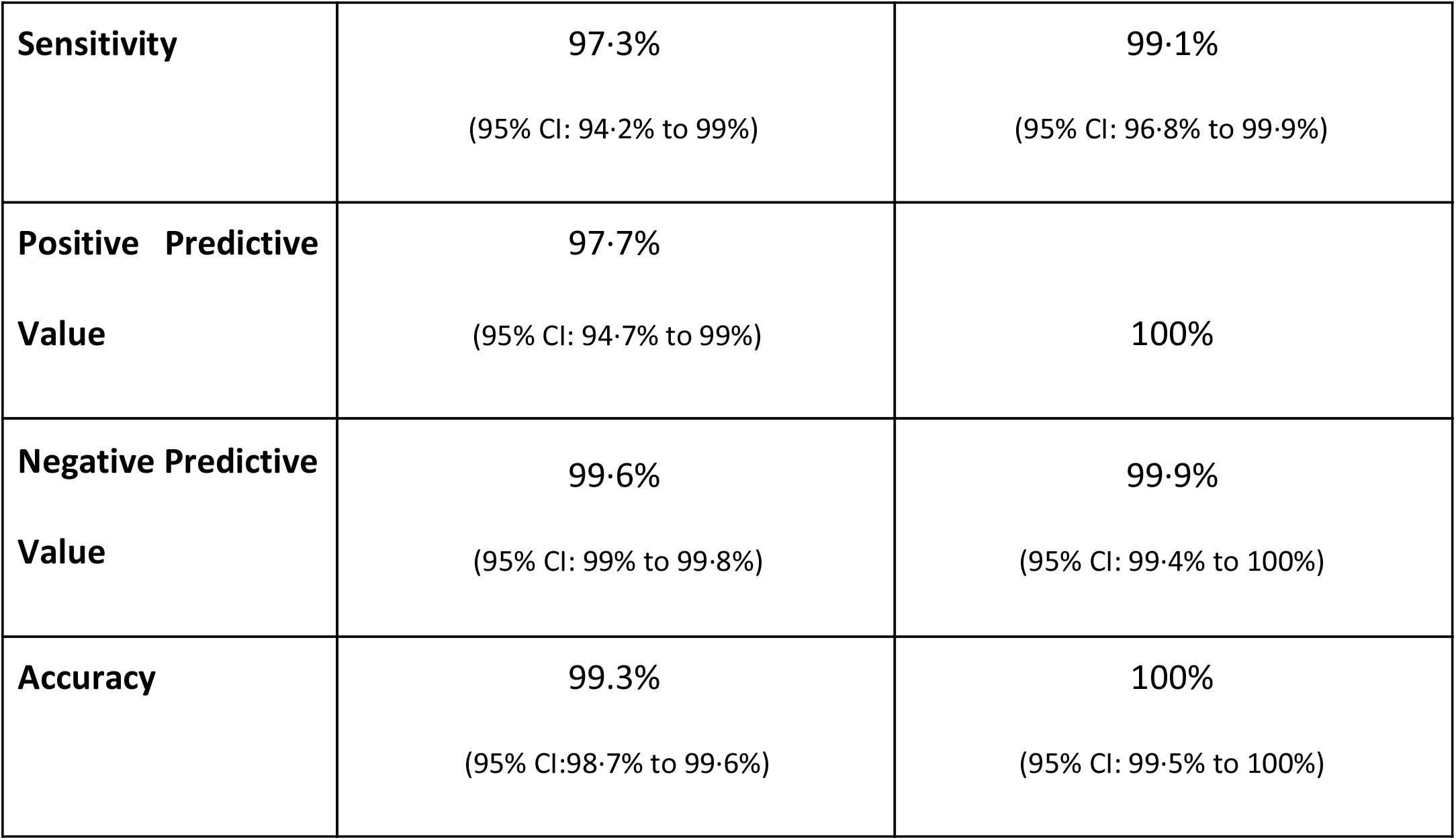
WGS repeat expansion detection performance. Performance based on total number of normal and expanded alleles across all loci tested after visual inspection. TN = True Negative; FP = False Positive; TP = True Positive; FN = False negative

|  | EHv3.1.2 |  |  |  | EHv312 after visual QC |  |  |  |
| --- | --- | --- | --- | --- | --- | --- | --- | --- |
|  | TN | FP | TP | FN | TN | FP | TP | FN |
| Total alleles | 1316 | 5 | 215 | 6 | 1321 | 0 | 219 | 2 |
| Specificity | 99.6%<br>(95% CI: 99.1% to 99.9%) |  |  |  | 100%<br>(95% CI: 99.7% to 100%) |  |  |  |

|  |  |  |
| --- | --- | --- |
| **Sensitivity** | **97·3%**<br>(95% CI: 94·2% to 99%) | **99·1%**<br>(95% CI: 96·8% to 99·9%) |
| **Positive Predictive Value** | **97·7%**<br>(95% CI: 94·7% to 99%) | **100%** |
| **Negative Predictive Value** | **99·6%**<br>(95% CI: 99% to 99·8%) | **99·9%**<br>(95% CI: 99·4% to 100%) |
| **Accuracy** | **99·3%**<br>(95% CI: 98·7% to 99·6%) | **100%**<br>(95% CI: 99·5% to 100%) |

As a further assessment of STR calling from WGS, repeat size estimates were compared with PCR-quantified lengths (**Table S4**). This dataset consisted of 418 PCR tests interrogating 805 alleles, 98% of which were normal and pre-mutation range (smaller than the WGS read-length (125bp or 150 bp) (see **Supplementary methods** for further details). 745 alleles sizes predicted by EH agreed with the PCR-assessed repeat lengths, yielding an overall concordance of 92.5%. Locus variability was observed, with higher concordance for *CACNA1A*, *ATXN2,* and *HTT* and lower for *TBP* (**Figure 2B**, **Table S6**). These results are consistent with other studies showing a high concordance between WGS and PCR quantification of repeat lengths smaller than the sequencing read length.^12,13,16^

The benchmark dataset included large expansions in *FMR1*, *DMPK*, *C9orf72,* and *FXN* which can extend to up to 5 kb in size. EH was able to correctly identify expanded alleles at these loci (**Table S5**), although EH size estimates trended lower as repeat size increased within the pathogenic range (**Figure 2, Table S7)**, and this affected the ability to distinguish between large and small expansions in DMPK or between full-expansions and premutations in *FMR1* (**Table S7**).

### Detection of pathogenic repeat expansions in undiagnosed individuals

To incorporate WGS-based repeat expansion detection within the clinical diagnostic setting, we applied our EH-enabled pipeline to WGS 13,331 individuals with a suspected genetic neurological disorder recruited to the 100,000 Genomes Project dataset. These individuals were drawn from four cohorts: (A) patients with a neurodegenerative disorder, including early onset dementia (*C9orf72)*, amyotrophic lateral sclerosis (*C9orf72*, and *AR*), hereditary ataxia (*ATN1*, *ATXN1*, *ATXN2*, *ATXN3*, *ATXN7*, *CACNA1A*, *FXN,* and *TBP*), hereditary spastic paraplegia (*FXN*), Charcot-Marie-Tooth (*AR*) andand early onset or complex parkinsonism (*ATXN2*, *ATXN3*) due to their overlapping presentations (n=3,626; **Table 2**); (B) paediatric patients with intellectual disability (ID) and seizures, dystonia, ataxia, spastic paraplegia, optic neuropathy, retinopathy, white matter abnormalities, muscular, or hypotonia (n=2,576; **Table 2** and **Supplementary Methods**) which were assessed for ‘CAG’ expansions in *HTT*, *ATN1*, *ATXN2*, *ATXN3*, *CACNA1A*, and *ATXN7*, as these REs have been reported to cause rare paediatric diseases.^17,18^; (C) paediatric and adult patients presenting with intellectual disability, a neuromuscular phenotype or a combination of the two (n=7,592; **Table 2**) assessed for *DMPK* expansions; and (D) paediatric patients presenting with intellectual disability alone analysed for *FMR1* expansions (n=6,731; **Table 2**).

**Table 2.**
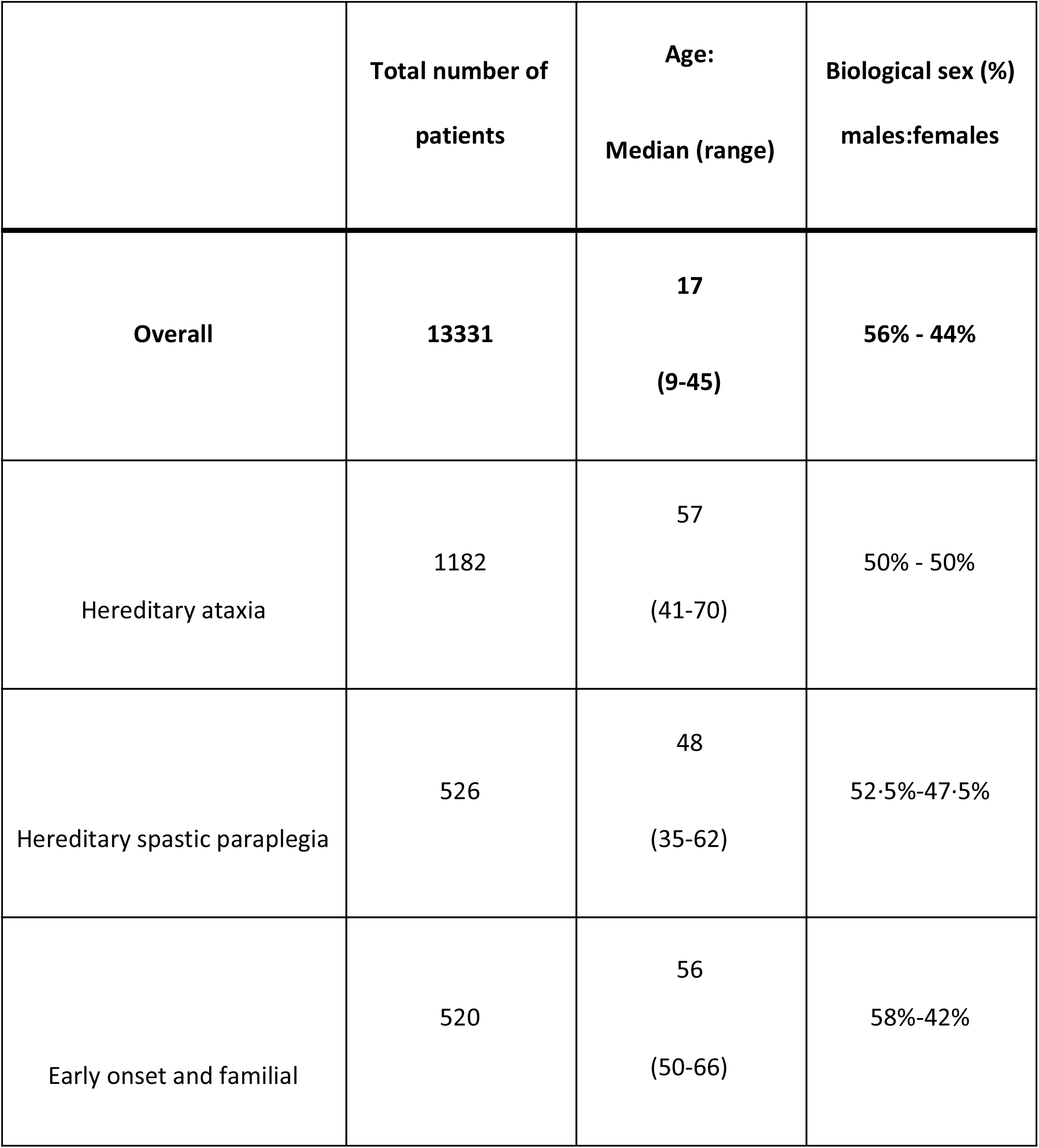
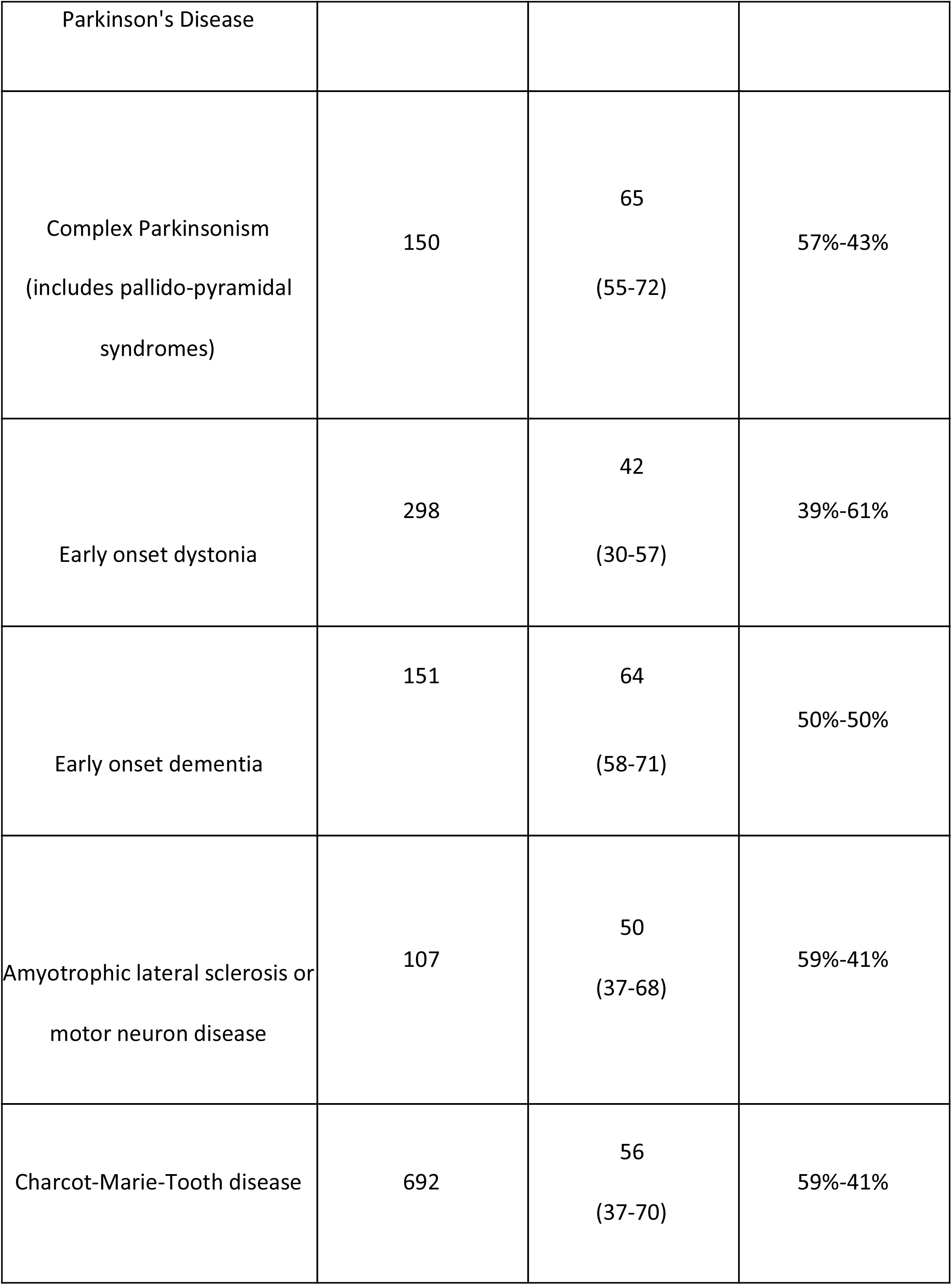
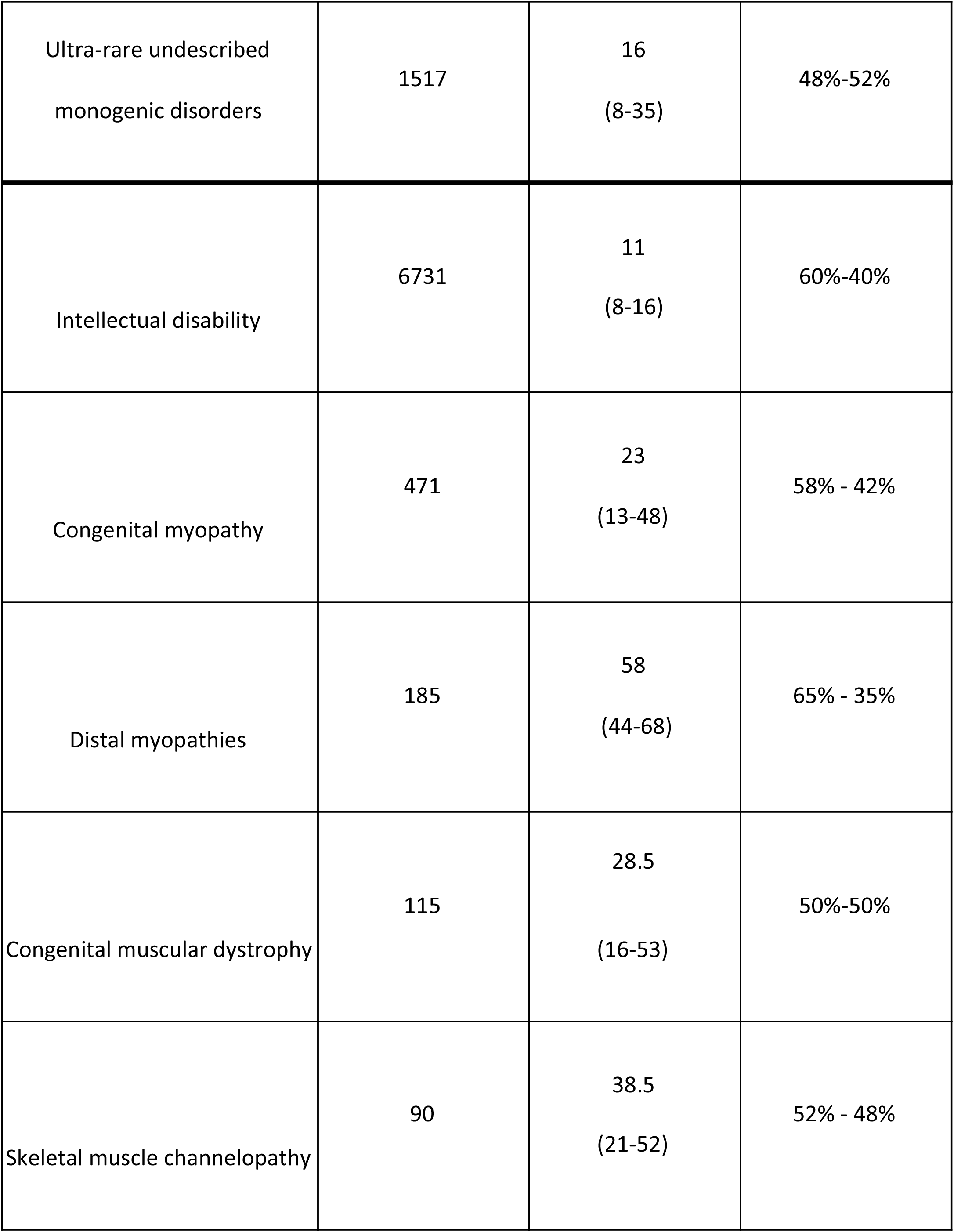
100,000 Genomes Project cohort analysed to identify pathogenic RE. Total number of patients, median age, and percentage of biological sex for each subsection. See **Table S14** for ancestry information.

Individuals who had previously tested negative after usual care testing in the NHS and participants with suspected monogenic disease and no prior molecular testing were eligible. Despite usual care NHS genomic testing being negative, we detected and visually confirmed repeat expansions in 96 individuals (**Table S8**). Of these, 82 cases were available for orthogonal testing, and 69 full expansions (**Table S9**). At the time of writing, diagnostic reports have been issued for 60 patients. Clinical details of the 69 individuals with full expansions are provided in **Table 3** and **Table S10**. Of the six EH calls that were not orthogonally confirmed, two were normal alleles in *ATXN1* and *ATXN2*, and four were *FMR1* intermediate range calls (n=4) (**Figure S5**). These results demonstrate that the RE aware WGS pipeline was successfully deployed at scale with a 0.09% false positive rate (13 false positive tests out of 13,331 individuals tested).

**Table 3.**
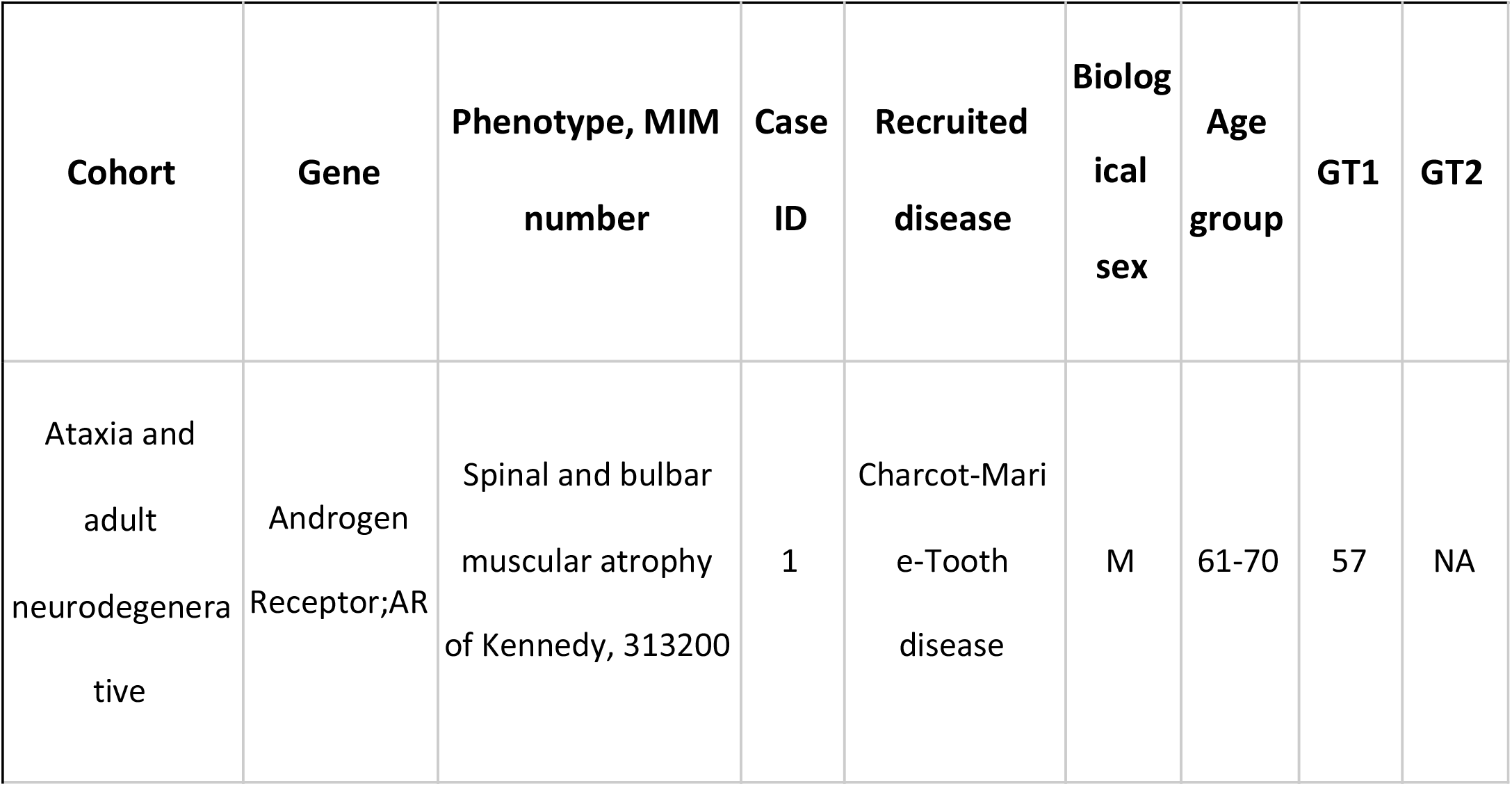
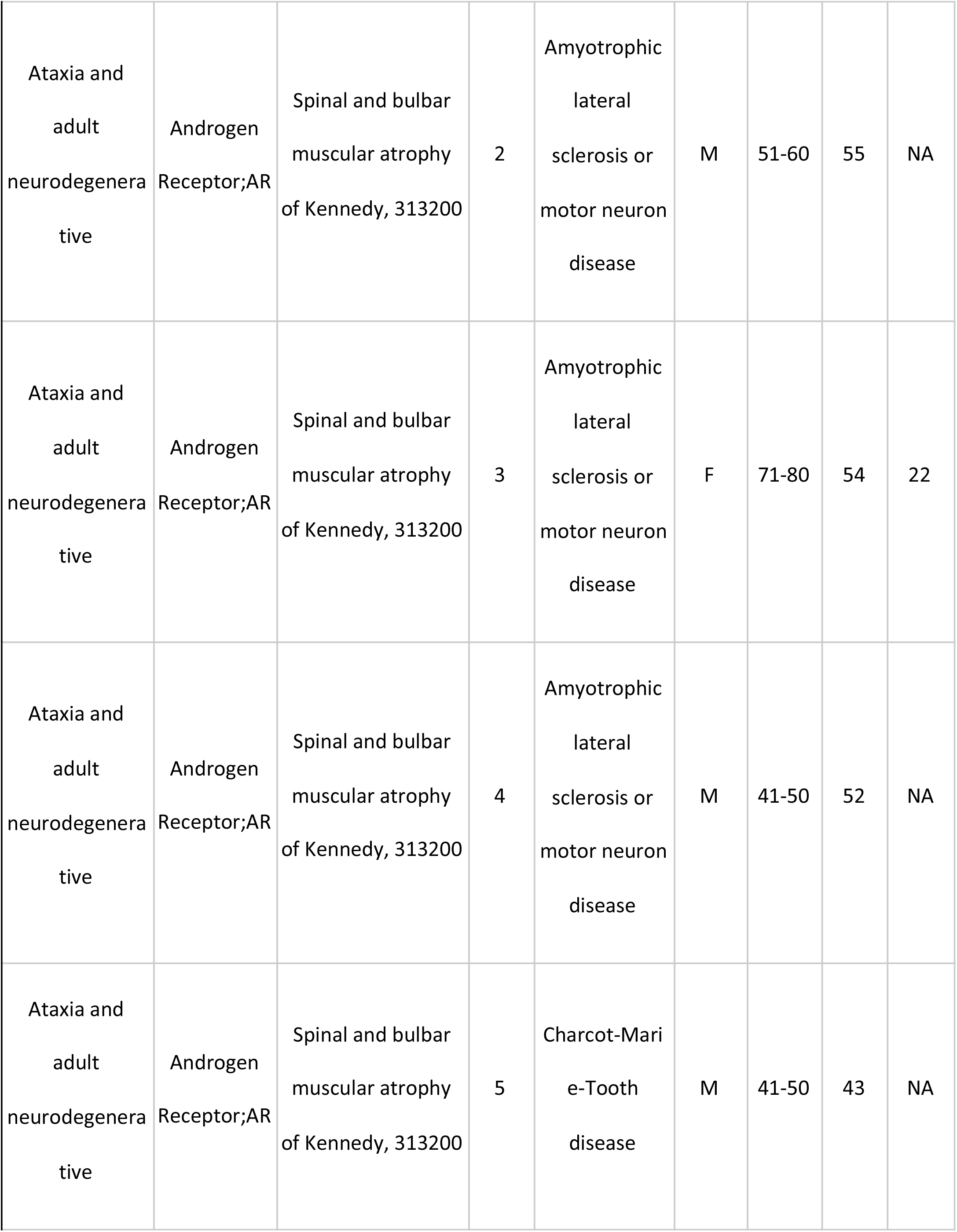
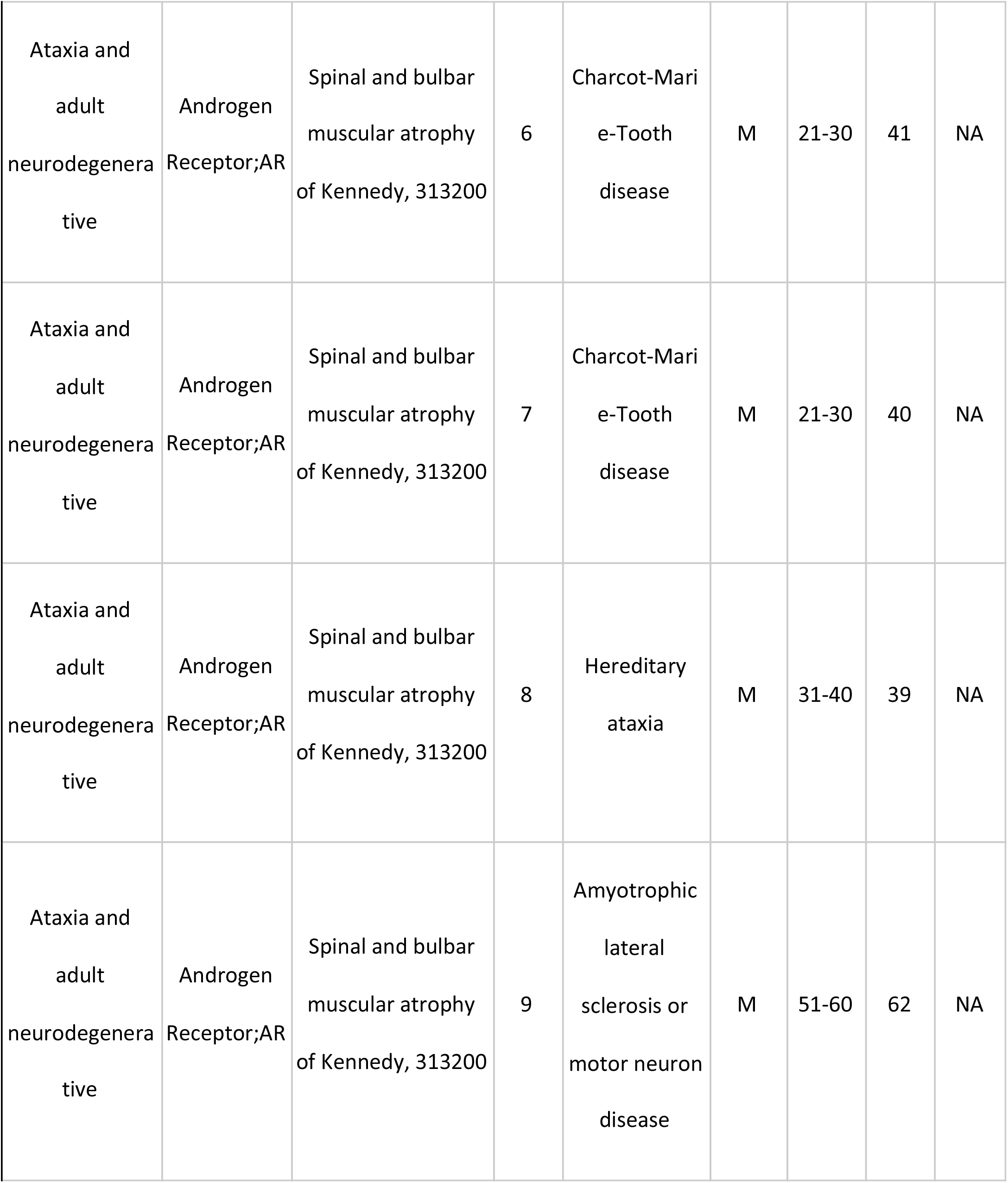
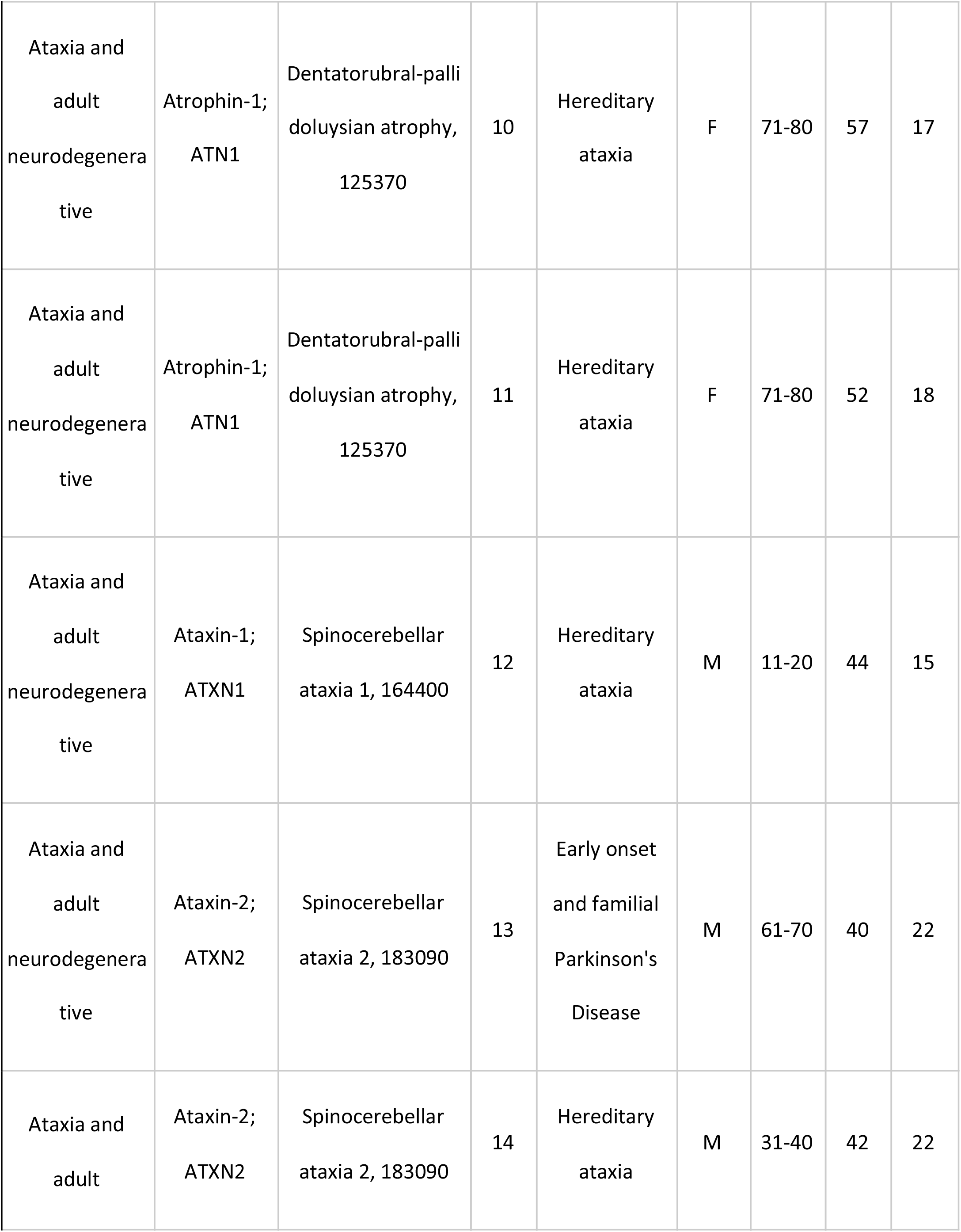
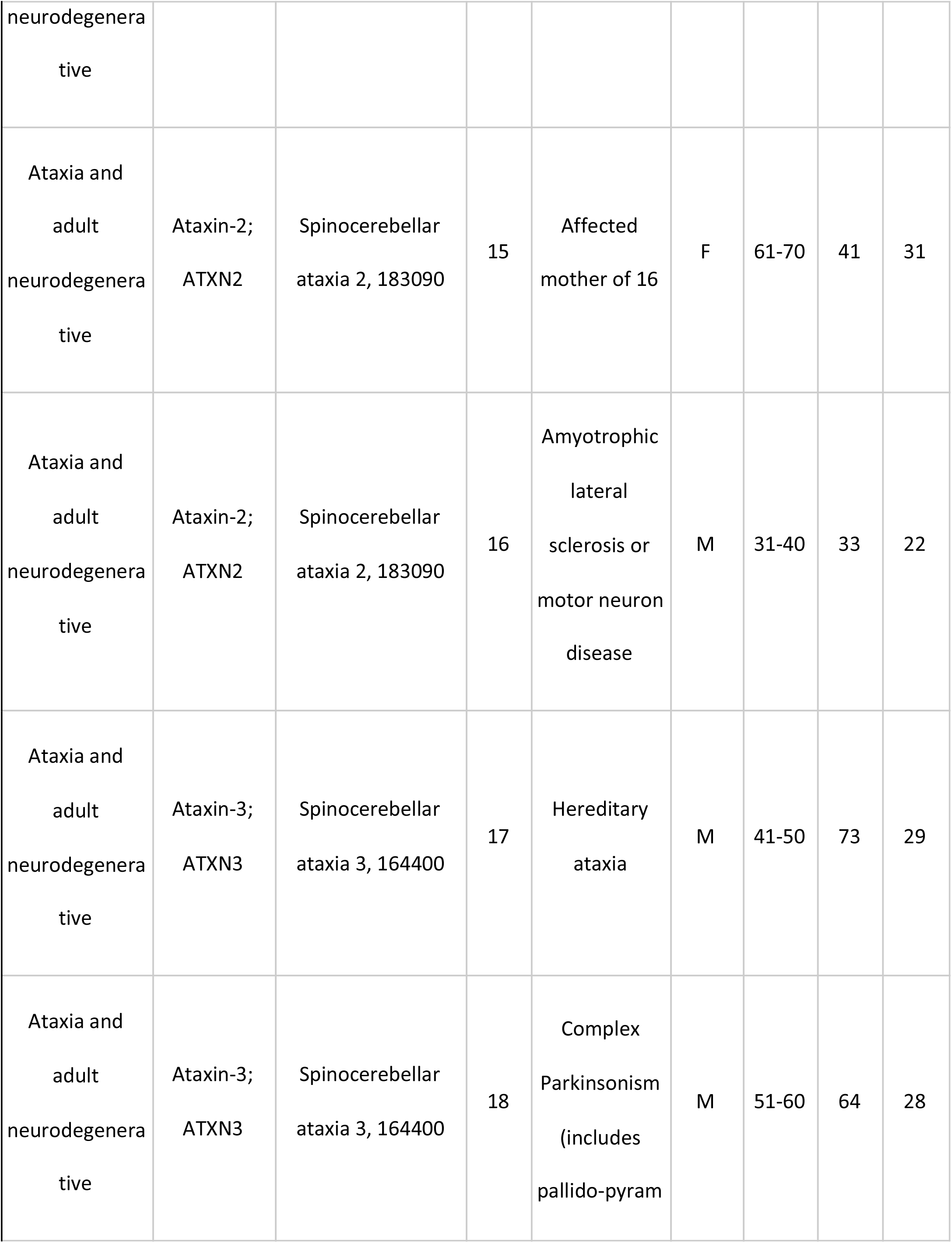
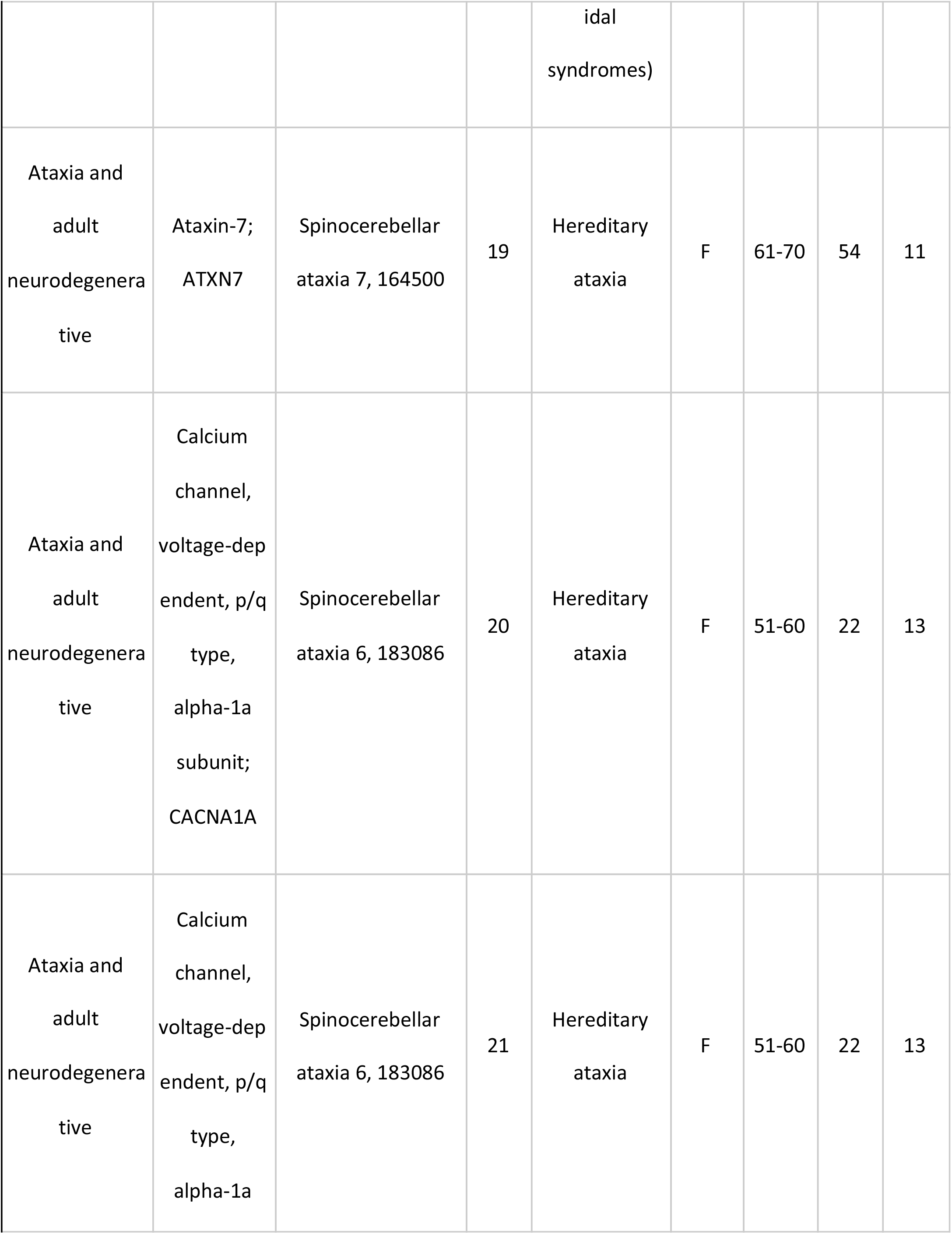
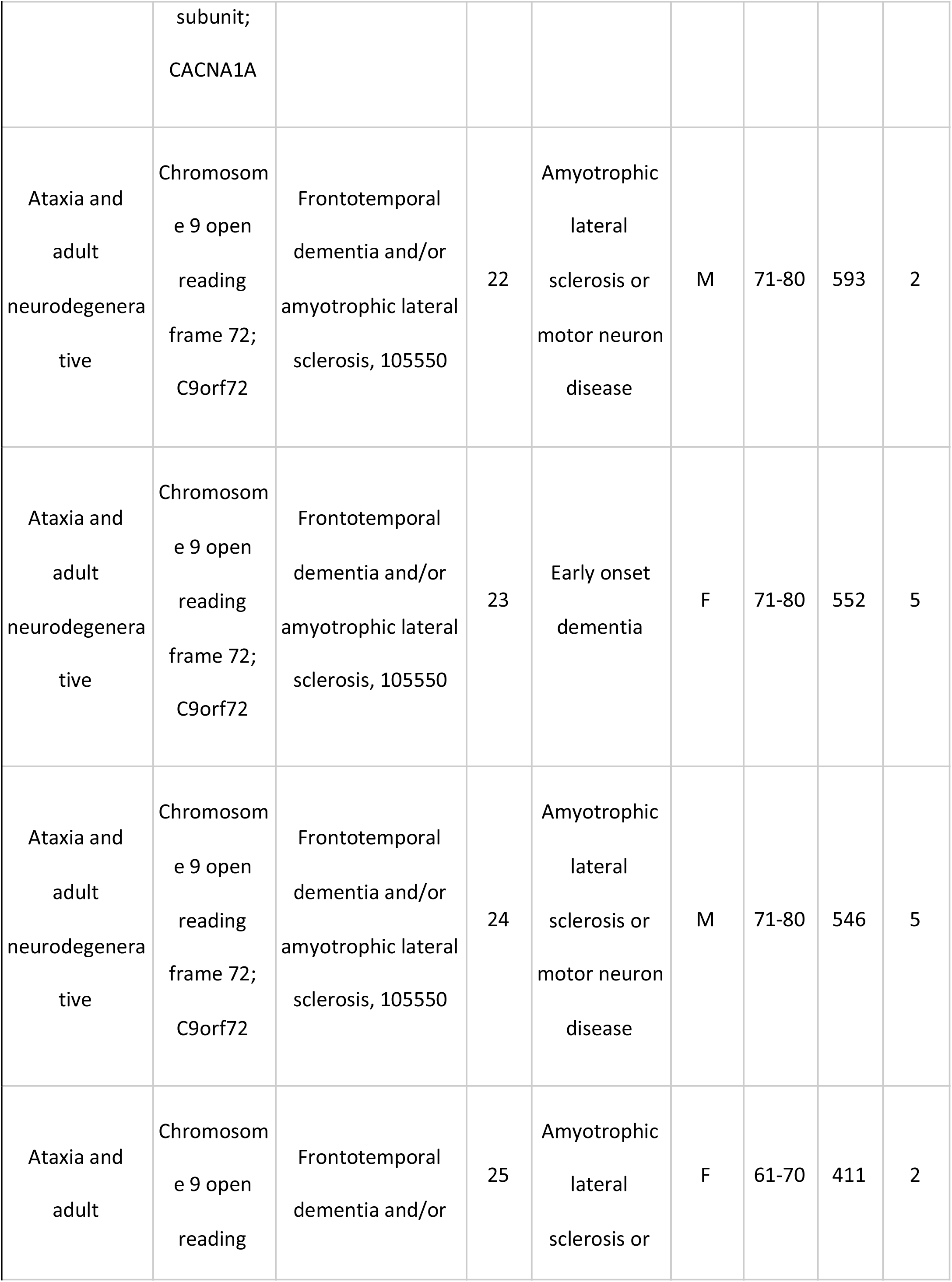
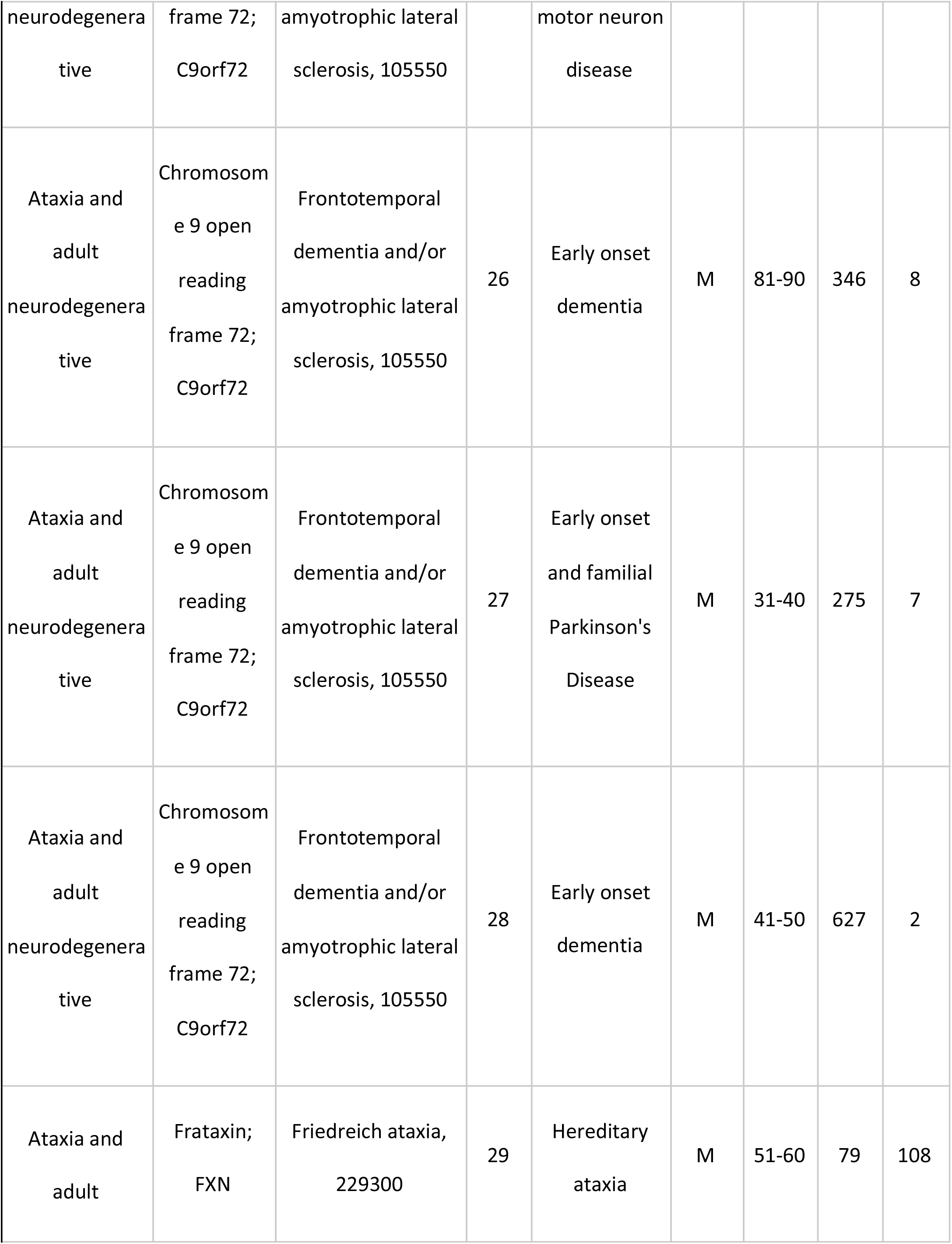
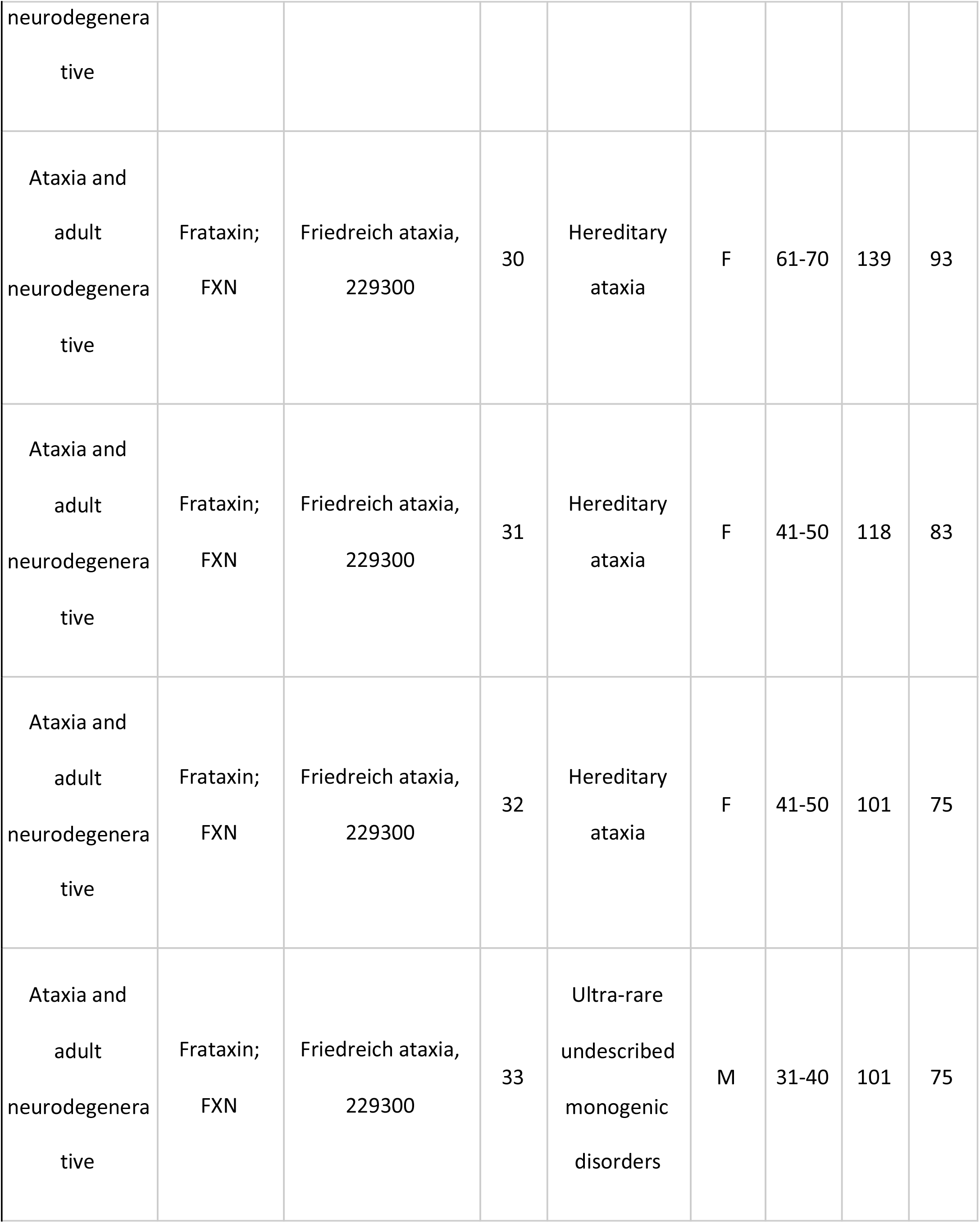
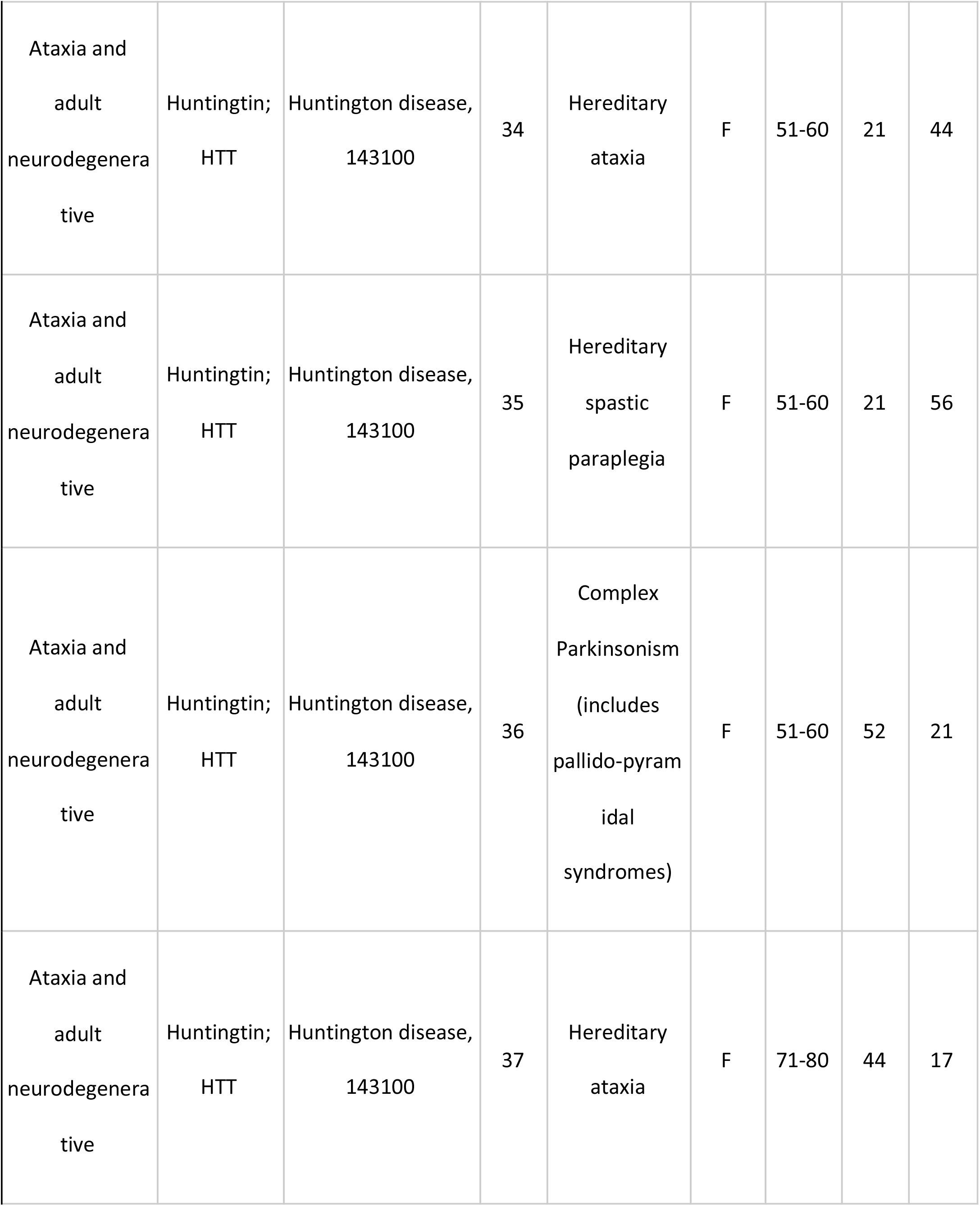
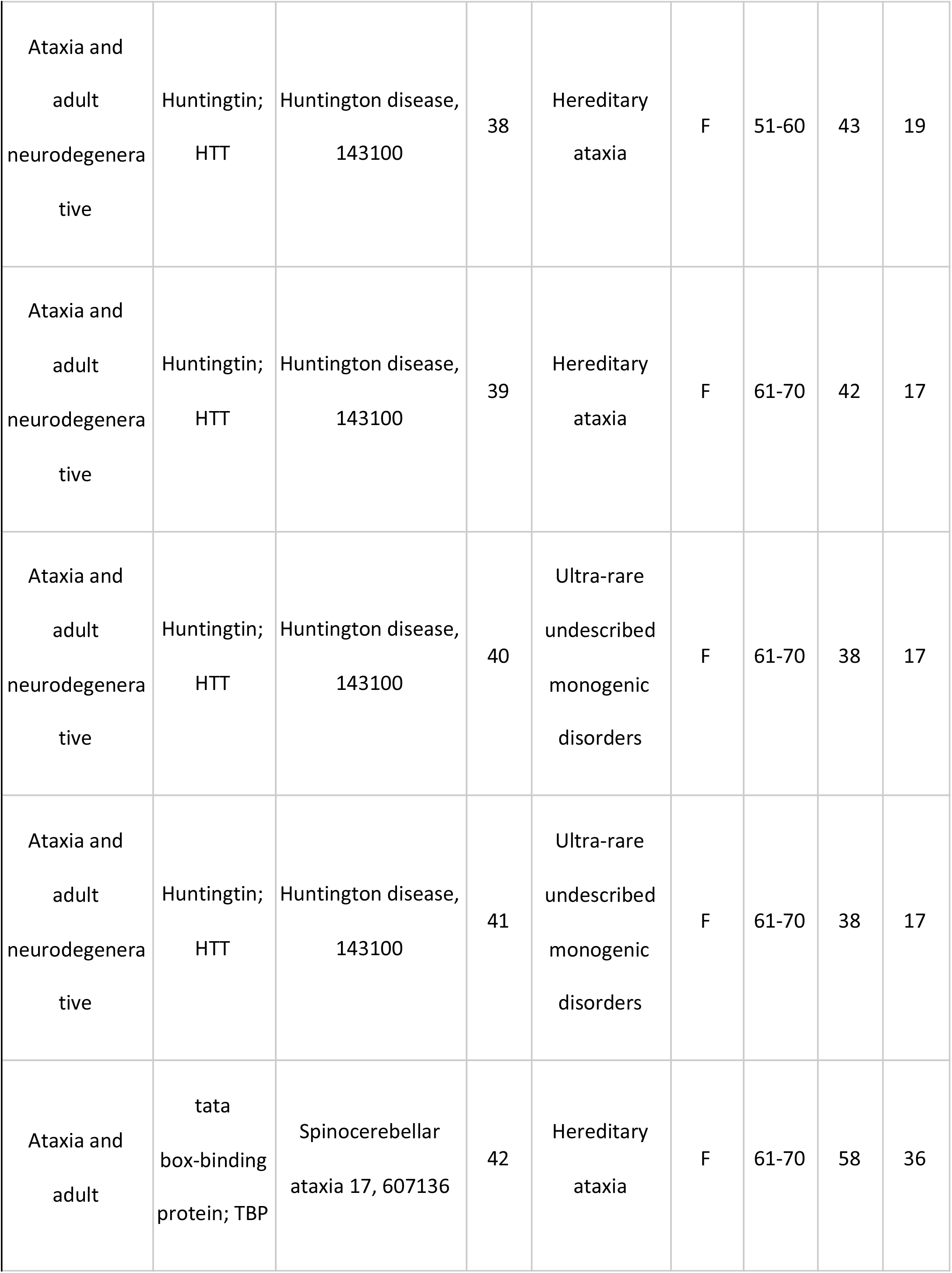
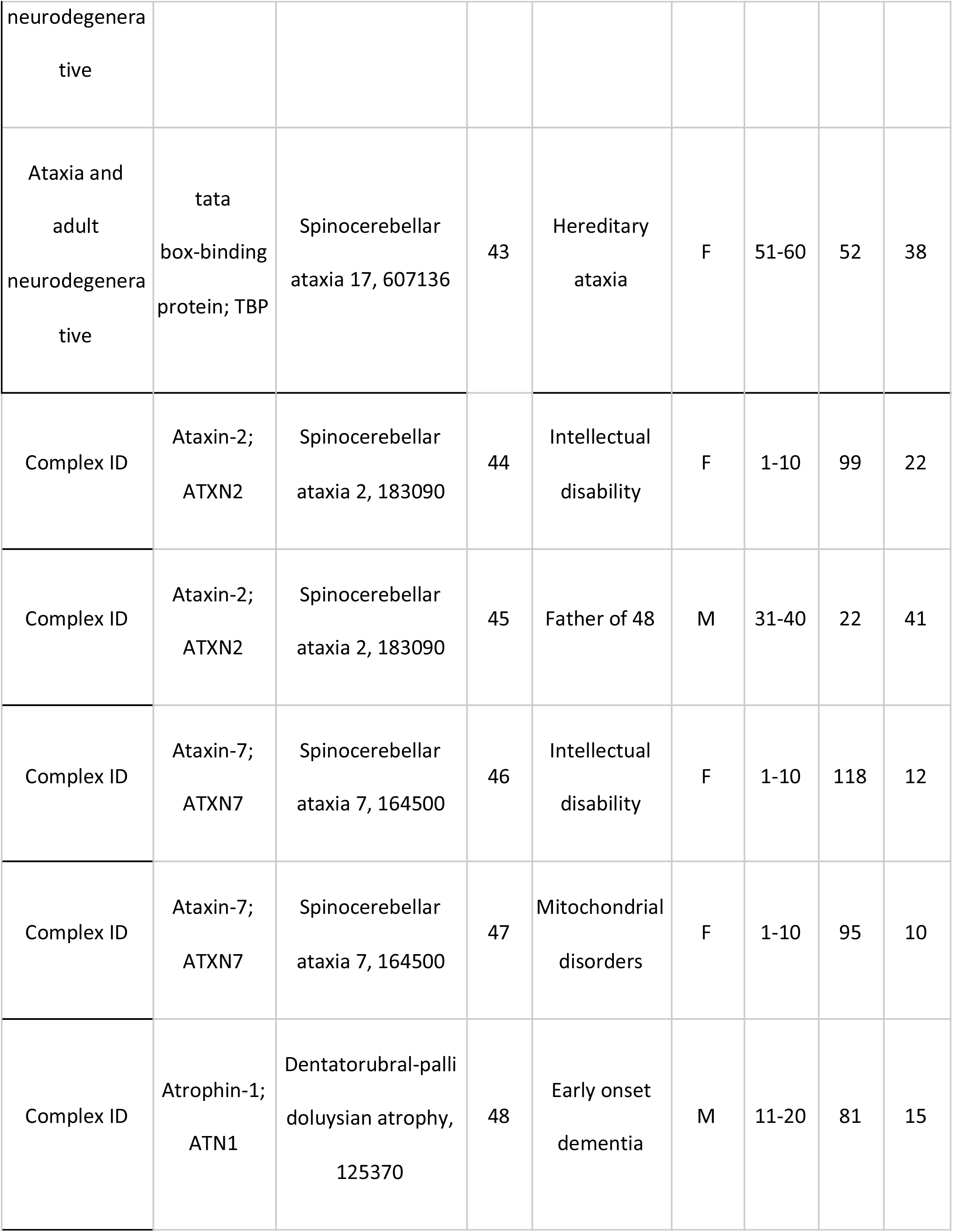
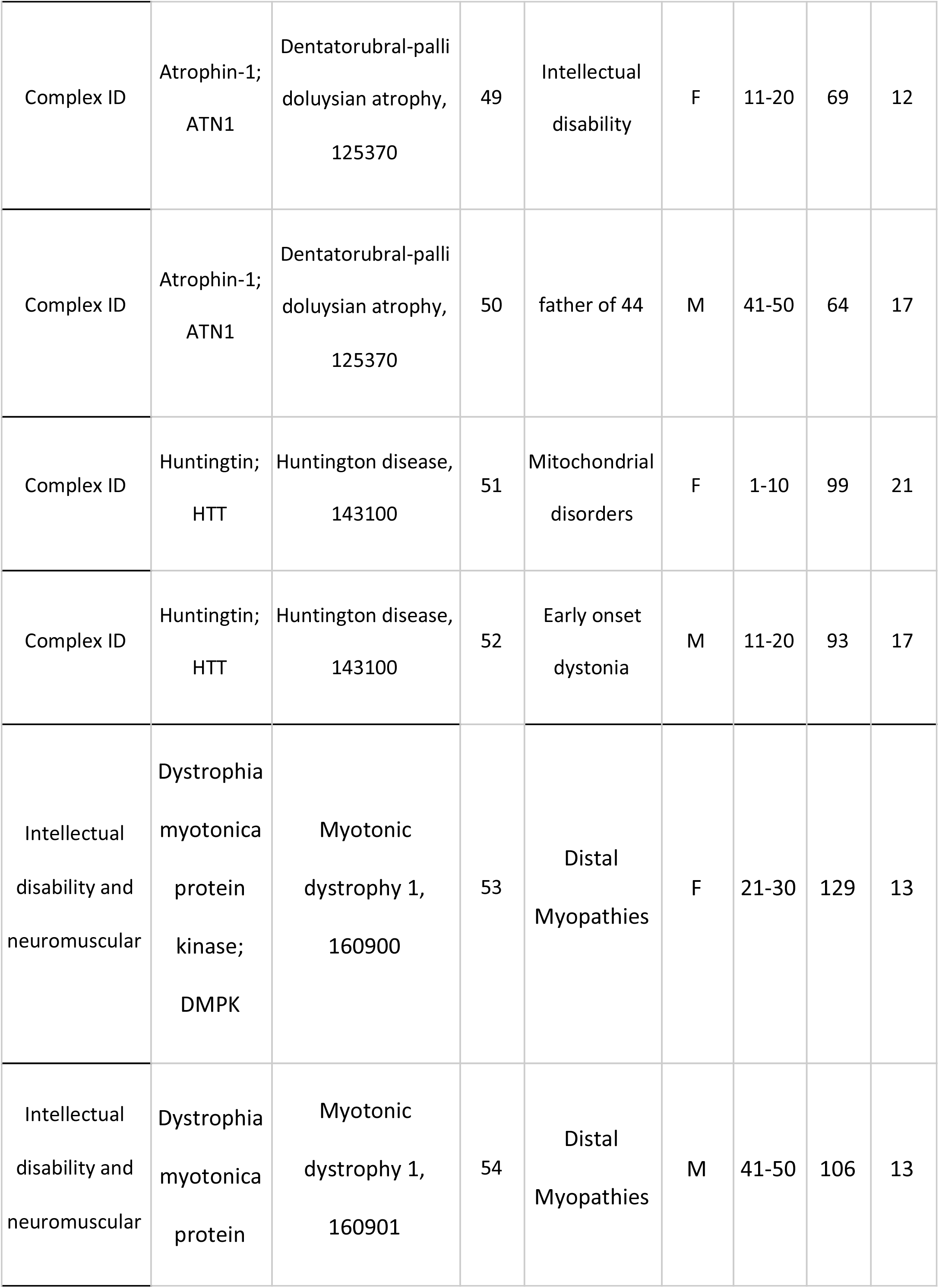
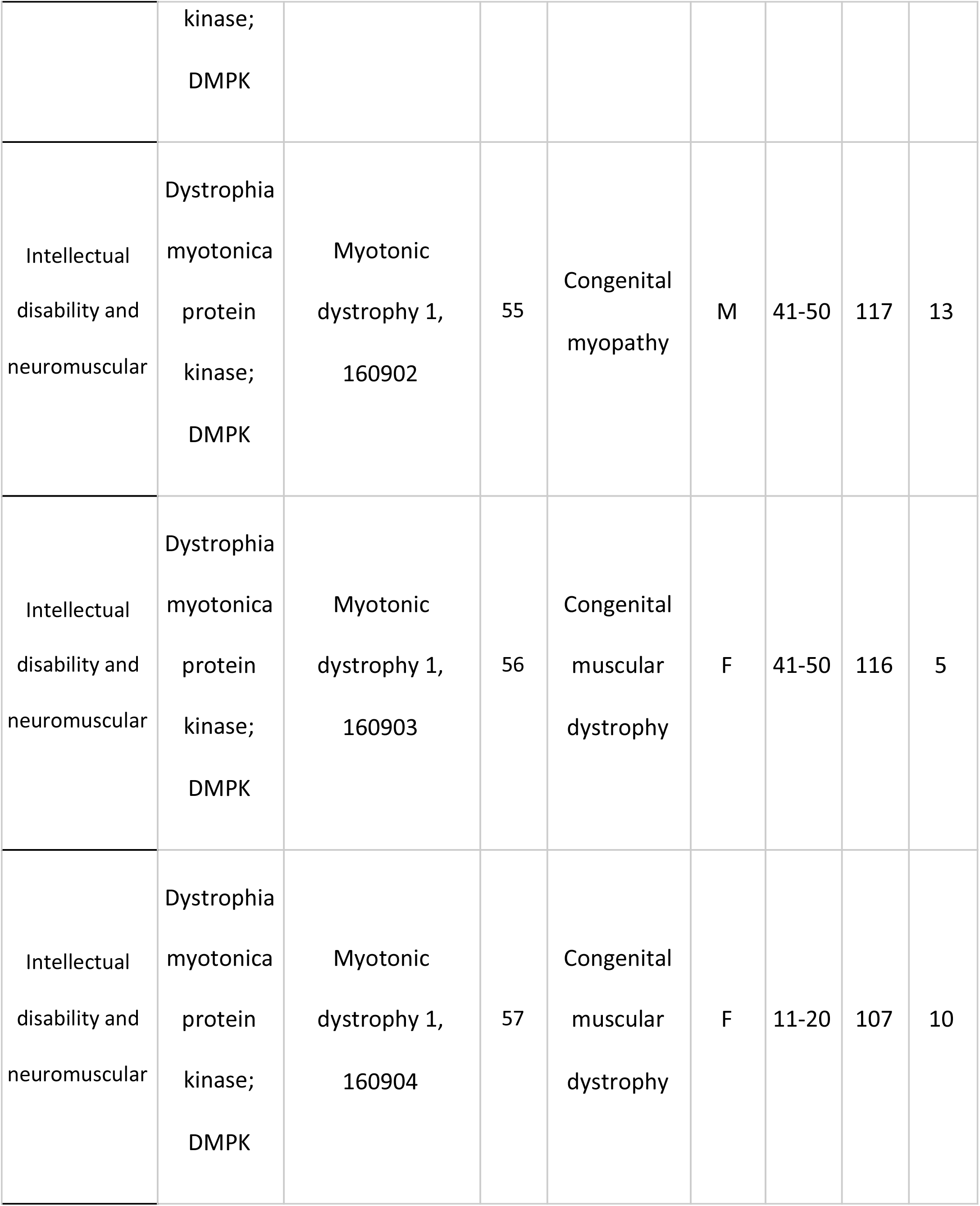
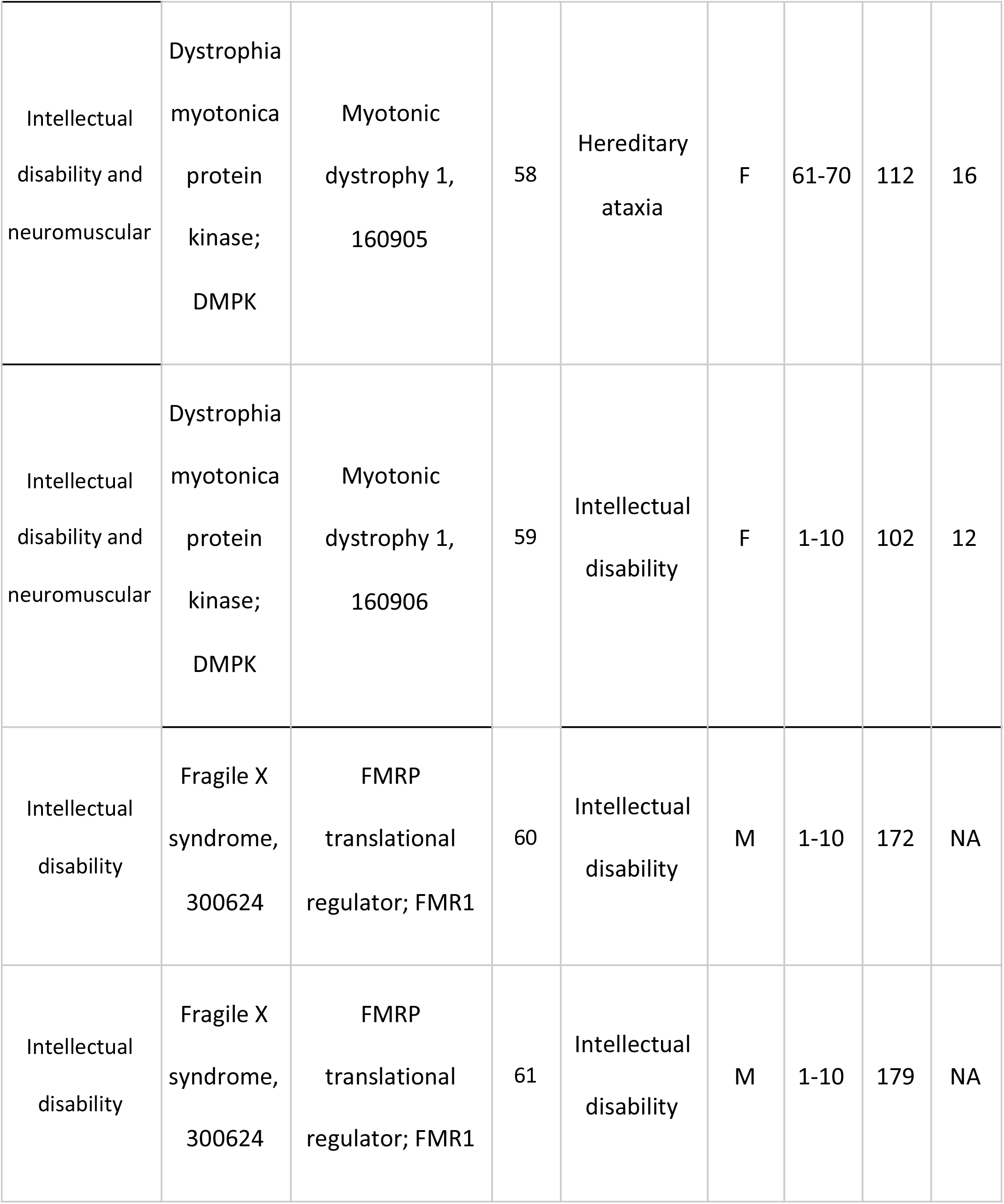
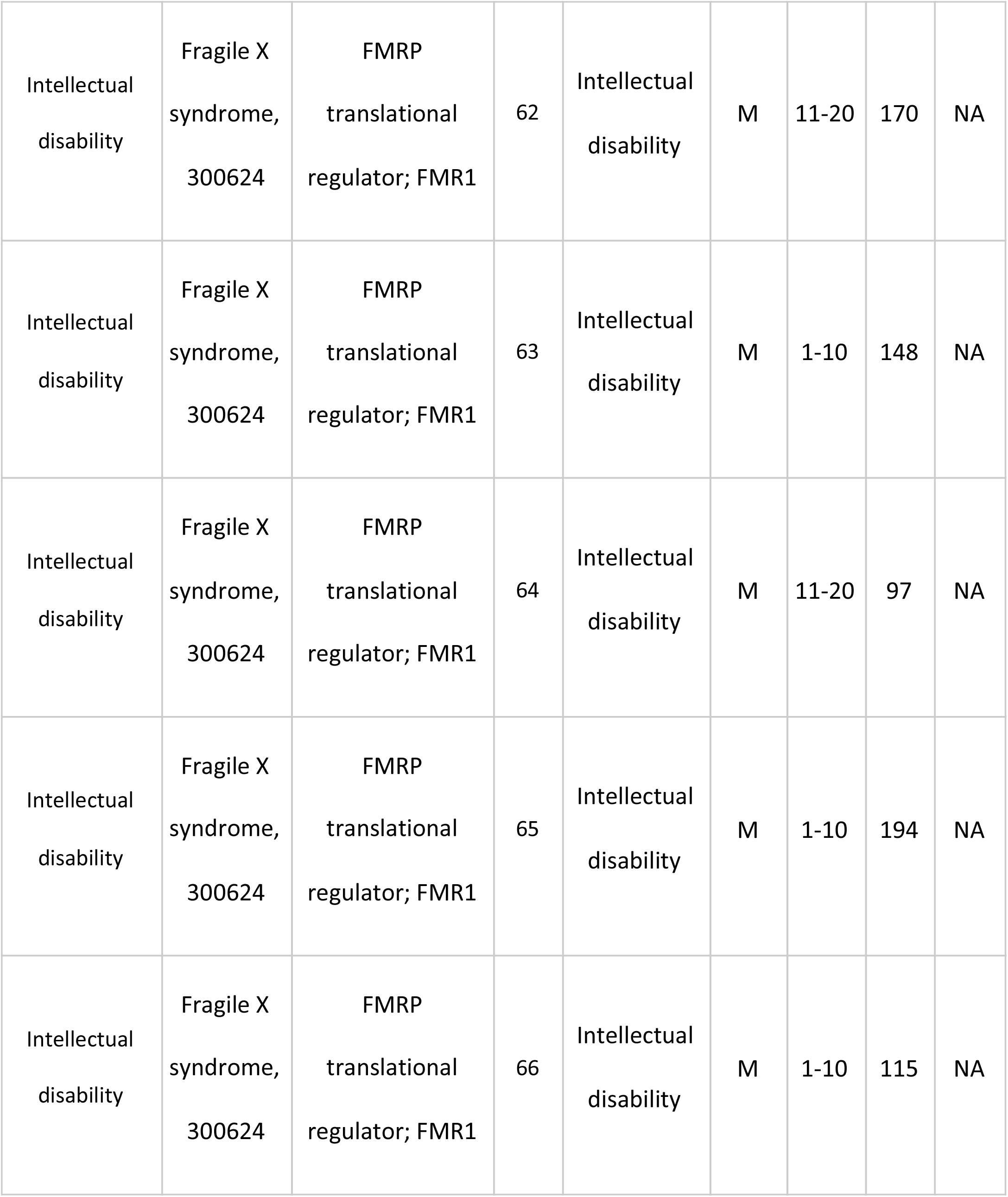
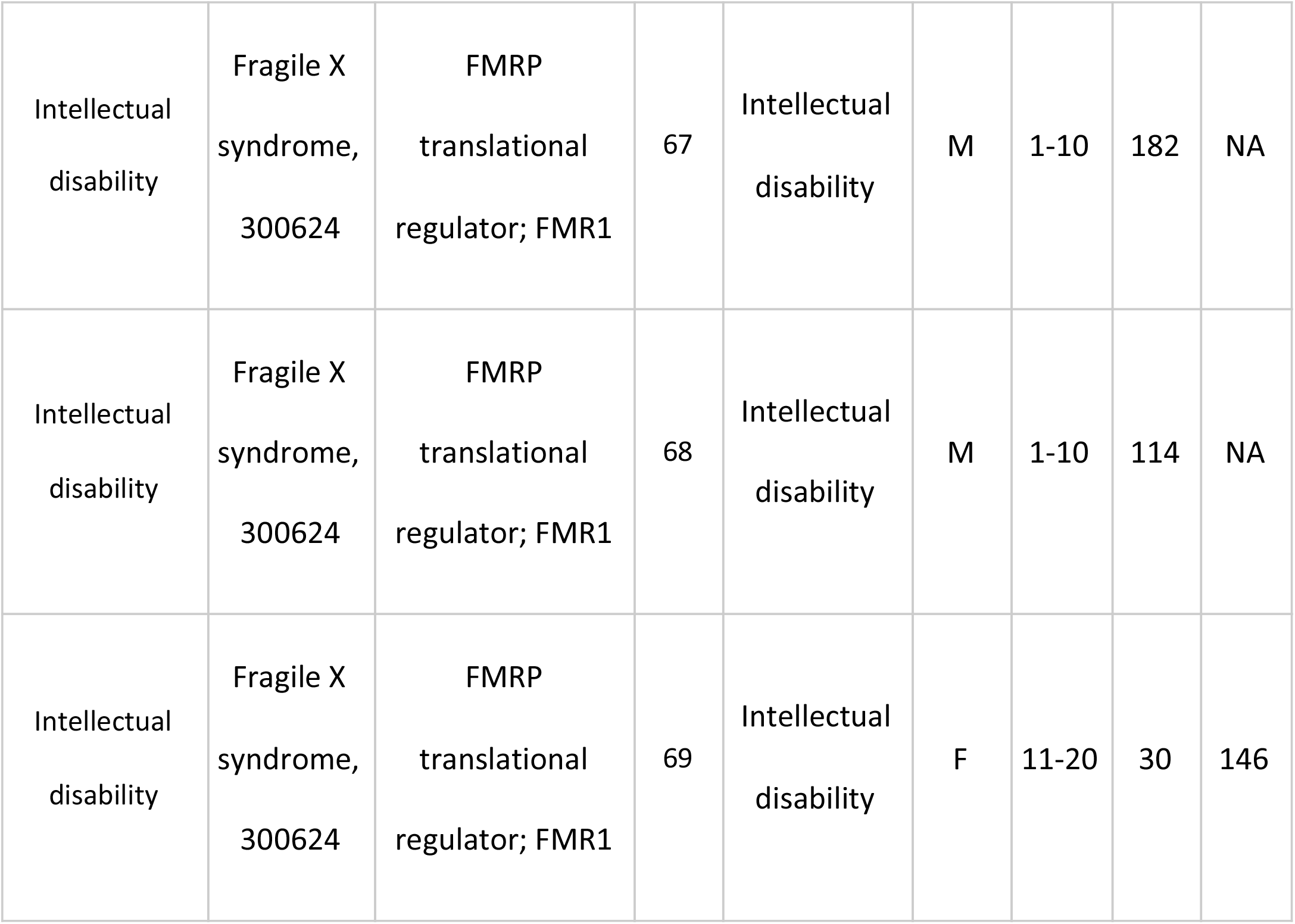
100,000 Genomes Project patients with pathogenic expansions. Patients with pathogenic expansions in the 100,000 Genomes Project. For each disease and gene (‘Gene’ and ‘Phenotype, ‘MIM number’ fields), information regarding the disease name under which a participant has been recruited together with biological sex, age range, and the genotypes are shown. The genotypes correspond to EHv2.5.5 sizes. The table is split into different disease groups (‘Cohort’). See **Table S10** for additional details including list of HPO terms for each individual.

In cohort A, expansions were observed in individuals presenting with a wide variety of overlapping phenotypes (**Table 3**), including an *ATXN2* RE in a individual with Levodopa-responsive early-onset Parkinson’s disease and a history of progressive cerebellar ataxia, an *AR* expansion in individuals clinically diagnosed with Charcot-Marie-Tooth disease including one with demyelinating neuropathy i.e. CMT1. Further, a wide range of prior clinical diagnoses were observed in individuals with pathogenic repeat expansions. For example, in seven individuals with amyotrophic lateral sclerosis or motor neuron disease, expansions were identified in *AR* (n=4) and *C9orf72* (n=3). In participants recruited under hereditary ataxia we identified expansions in loci that had not been assessed within the NHS at the time of recruitment, including *ATN1*, *ATXN2, ATXN3, ATXN7, CACNA1A, FXN, TBP* (**Table 3**). We also detected REs in individuals with a phenotype that was consistent with a different repeat expansion disorder, e.g. a *C9orf72* expansion in early onset and familial Parkinson’s Disease (case 27, **Table 3**), and repeat expansions in the reduced penetrance range (38 repeats in *HTT* in two sisters with movement disorder, dementia, depression and speech difficulties, cases 40 and 41 **Table S10**) underscoring the diagnostic challenge presented by these disorders. Taken together, these data demonstrate that the diagnosis challenges due the complexity of overlapping and pleiotropic presentation of repeat expansions disorders in adults can be reduced with a whole genome testing approach.

Strikingly, seven children in cohort B were identified with large ‘CAG’ expansions (**Figure 3**). Six lacked any informative family history and had not been offered RE testing as part of their clinical genomic testing at the time of recruitment (**Table 3**). Two children under the age of 10 carried large *HTT* expansions (90-100 ‘CAG’ repeats). Remarkably, one child had inherited the repeat from an unaffected parent with no family history of Huntington disease. Family testing is ongoing, but a reduced penetrance allele has been identified in the wider family, indicating that the repeat had expanded by over 60 repeat units in a single generation (case 52, **Table 3**). Two children under the age of five carried large *ATXN7* expansions and presented with complex multi-system phenotypes. This included a girl (case 46, **Table 3**), whose parent began to show gait problems two years after enrolment in the 100,000 Genomes Project. Similarly, a ten year old girl with an indication for testing of intellectual disability was found to have a *ATXN2* expansion of 99 repeats, despite both parents recruited to the project being designated as ‘unaffected’, and a 18 year old girl with dementia was found to carry an 69 repeat expansion in *ATN1* (**Table 3**). These data suggest that genome-wide testing of repeat loci can resolve cryptic pediatric genetic disease cases.

**Figure 3.**
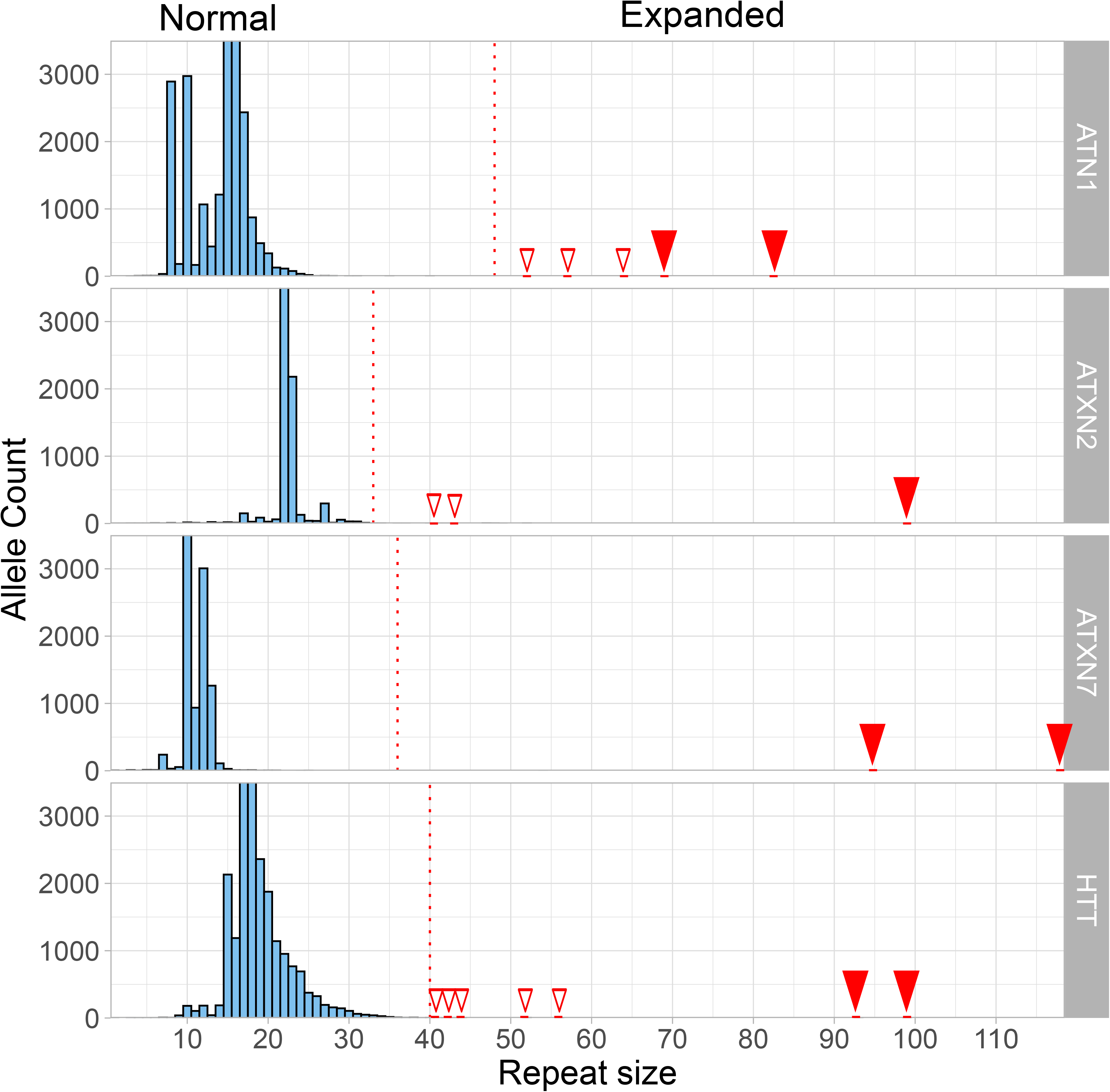
Adult and paediatric cases showing pathogenic expanded repeats. Repeat size frequency distribution of *ATN1* **(A)**, *ATXN2* **(B)**, *ATXN7* **(C),** and *HTT* **(D)** in 13,331 individuals; ‘CAG’ repeat count in X axis; allele count in Y axis. The dotted red line represents the full mutation threshold for each locus (**Table S3**). White and red arrowheads indicate adult and paediatric pathogenic expansions respectively.

In cohort C, seven expansions in *DMPK* were confirmed in five families, including a child and a mother with a clinical diagnosis of muscular dystrophy, two siblings with a suspected ‘distal myopathy’ disease (cases 53 and 54 **Table 3**), and one case with a suspected ataxia that also presented with muscular weakness (case 58). Lastly, in cohort D, *FMR1* expansions were detected in seven males, where a diagnosis of Fragile X syndrome fully or partially explained the presenting phenotype (cases 60-66, **Table 3**).

## DISCUSSION

The diagnosis of RE disorders is challenging in healthcare due to heterogeneous clinical presentations, overlapping phenotypes and non-specific clinical findings, which may increase in severity with age, and in each subsequent generation. Repeat expansion disorders are amongst the most common causes of inherited neurological diseases.^2^ Nonetheless, patients may be underdiagnosed because of the fragmented testing approaches currently employed - they may have the incorrect repeat expansion locus tested^19^ or receive a molecular test for a different class of variant due to the phenotypic overlap with other neurological genetic disorders.^20^

WGS has been deployed in multiple settings as a first line diagnostic test for rare neurological disorders, but has previously been thought to have limited ability to detect repeat expansions.^11^ Recently, several tools have been developed to call repeats from WGS in the research setting.^21^ However, none has been implemented in a clinical setting. In this study, we present evidence that a WGS bioinformatic pipeline incorporating an accurate expansion-aware algorithm can reliably assess the most common disease-causing repeat expansions and resolve previously intractable cases in a large cohort of patients with genetically undiagnosed neurological disorders.

When WGS RE detection was assessed against positive and negative controls previously characterized in clinical diagnostic genomic laboratories using gold standard methods, we found an overall minimum 97.3·6% sensitivity and 99·6% specificity. This reflects the ability of the pipeline to accurately discriminate between normal and disease-causing alleles across 13 RE disorder loci. Furthermore, these data show that repeat sizing is accurate for repeats smaller than the sequencing read lengths, and therefore most normal and premutation alleles for the ‘CAG’ repeat expansion disorders can be sized accurately. The WGS expansion detection pipeline is limited in its sizing of alleles significantly larger than the read-length, such as Fragile X. For example, we note that all *FMR1* repeats previously classified by PCR as ‘expanded’ were classified by WGS as premutation in this study.

Our findings enable the establishment of a clinical diagnostic workflow for WGS (**Figure S6**) in which repeat expansions are classified as either ‘normal’ or ‘expanded’ (i.e. larger than the premutation cut-off) without adherence to the repeat size estimation, particularly for large repeats. We propose that all WGS-calls classified as ‘expanded’ are visually inspected to detect false positive calls (**Figure S2**) and detect the presence of biallelic expansions where only one expanded allele has been detected (e.g. *FXN*) (cases A-C, **Figure S3**). Additionally, we recommend that after visual confirmation the presence of the expansion is validated by orthogonal testing.

The application of the WGS-pipeline to undiagnosed patients tested in the ICSL laboratory has led to five RE diagnoses (in *ATXN2*, *FXN* & *DMPK*), including the detection of pathogenic mosaic maternal expansions. In participants recruited to the 100,000 Genomes Project, pathogenic REs consistent with the patient’s phenotypes have been diagnostically confirmed in 60 cases. Remarkably, some of the expansions were not suspected based on the patient phenotype, including six paediatric subjects without any family history of a RE disorder. The average repeat expansion sizes detected across the paediatric cases described here are substantially larger than the average in adults, strongly suggesting that using age-specific repeat-size thresholds may eliminate any potential hazard of identifying adult-onset risk alleles in children (**Figure 3**).

Rare inherited diseases may present with a wide phenotypic spectrum that often overlaps multiple different syndromes making locus specific genomic testing inefficient, arduous, and expensive. We present evidence here that a clinical grade WGS bioinformatic pipeline, with potential to diagnose a range of rare neurologic diseases, may now be extended to identify REs. Since WGS provides a single test that can identify the most common REs, it offers the opportunity to identify the majority of patients with these heterogeneous disabling disorders where the diagnosis may be missed by locus-specific testing. In the era of emerging therapies for these disorders early detection may become crucial.^22^ As a result this is now being considered for adoption in the NHS England National Genomic Test Directory for application to undiagnosed rare neurologic disease in direct healthcare.

## Acknowledgements

Genomics England and the 100,000 Genomes Project was funded by the National Institute for Health Research, the Wellcome Trust, the Medical Research Council, Cancer Research UK, the Department of Health and Social Care and NHS England. We thank all the participants and healthcare teams at the 13 NHS Genomic Medicine Centres where ~5000 multidisciplinary staff enrolled patients to the 100,000 Genomes Project in the East of England, Greater Manchester, North East and North Cumbria, North Thames, North West Coast, Oxford, South London, West London, West Midlands, South West, Wessex, West of England and Yorkshire and Humber. Participants were also enrolled from Scotland by the Scottish Genomes Project, across Wales and Northern Ireland.

This work forms part of the portfolio of translational research at the NIHR Biomedical Research Centres at Barts, Birmingham, Bristol, Cambridge, Great Ormond Street Foundation, Guy’s and St Thomas’s, Imperial, Leeds, Leicester, Manchester, Maudsley, Moorfields, Newcastle, Nottingham, Oxford, Royal Marsden, Sheffield, Southampton and University College London. This work was made possible through the generosity of NHS patients and their families and uses clinical data from the NHS and NHS Digital. The views expressed are those of the author(s) and not necessarily those of the NHS, the NIHR or the Department of Health and Social Care.

Mark Caulfield is an NIHR Senior Investigator. Patrick Chinnery is a Wellcome Trust Principal Research Fellow (212219/Z/18/Z), and an NIHR Senior Investigator, who receives support from the Medical Research Council Mitochondrial Biology Unit (MC_UU_00015/9), the Medical Research Council (MRC) International Centre for Genomic Medicine in Neuromuscular Disease (MR/S005021/1), the Leverhulme Trust (RPG-2018-408), an MRC research grant (MR/S035699/1), an Alzheimer’s Society Project Grant (AS-PG-18b-022). John Sayer is supported by Kidney Research UK (RP_006_20180227). Richard Festenstein receives support from Imperial NIHR BRC. Jonathan M Schott receives support from National Institute for Health Research University College London Hospitals Biomedical Research Centre, Brain Research UK (UCC14191) and Medical Research Council. Lyn S Chitty is an NIHR Senior Investigator and is partly funded by GOSH NIHR Biomedical Research Centre, which has provided infrastructure support for the North Thames GMC. Arianna Tucci is an MRC Clinician Scientist (MR/S006753/1).

We thank Illumina Clinical Laboratory Sciences, San Diego and Illumina, San Diego and Granta Park Cambridge for undertaking whole genome sequencing and validation of true positives and negatives. We are grateful for the support of Dom McMullan, Helen Firth, Steve Abbs, Sian Ellard for their role in supporting the development of the bioinformatics pipeline and reporting process. We are grateful for the input and support from Professor Dame Sue Hill and the team in NHS England and for the work to fund and establish the 13 Genomic Medicine Centres. This enabled the NHS contribution to the 100,000 Genomes Project by enrolment of patients, receipt of the results and in some cases orthogonal validation using standardised approaches including return of findings for direct patient benefit.

