## Supplementary Methods for "Whole genome sequencing for diagnosis of neurological repeat expansion disorders"

#### INVESTIGATORS

The members of The Genomics England Research Consortium are:

J. C. Ambrose[1], P. Arumugam[1], E. L. Baple[1], M. Bleda[1], F. Boardman-Pretty[1,2], J. M. Boissiere[1], C. R. Boustred[1], H. Brittain[1], M. J. Caulfield[1,2], G. C. Chan[1], C. E. H. Craig[1], L. C. Daugherty[1], A. de Burca[1], A. Devereau[1], G. Elgar[1,2], R. E. Foulger[1], T. Fowler[1], P. Furió-Tarí[1], J. M. Hackett[1], D. Halai[1], A. Hamblin[1], S. Henderson[1,2], J. E. Holman[1], T. J. P. Hubbard[1], K. Ibáñez[1,2], R. Jackson[1], L. J. Jones[1,2], D. Kasperaviciute[1,2], M. Kayikci[1], L. Lahnstein[1], K. Lawson[1], S. E. A. Leigh[1], I. U. S. Leong[1], F. J. Lopez[1], F. Maleady-Crowe[1], J. Mason[1], E. M. McDonagh[1,2], L. Moutsianas[1,2], M. Mueller[1,2], N. Murugaesu[1], A. C. Need[1,2], C. A. Odhams[1], C. Patch[1,2], D. Perez-Gil[1], D. Polychronopoulos[1], J. Pullinger[1], T. Rahim[1], A. Rendon[1], P. Riesgo-Ferreiro[1], T. Rogers[1], M. Ryten[1], K. Savage[1], K. Sawant[1], R. H. Scott[1], A. Siddiq[1], A. Sieghart[1], D. Smedley[1,2], K. R. Smith[1,2], A. Sosinsky[1,2], W. Spooner[1], H. E. Stevens[1], A. Stuckey[1], R. Sultana[1], E. R. A. Thomas[1,2], S. R. Thompson[1], C. Tregidgo[1], A. Tucci[1,2], E. Walsh[1], S. A. Watters[1], M. J. Welland[1], E. Williams[1], K. Witkowska[1,2], S. M. Wood[1,2], M. Zarowiecki[1]

1. Genomics England, London, UK

2. William Harvey Research Institute, Queen Mary University of London, London, EC1M 6BQ, UK

### SUPPLEMENTARY METHODS

#### Repeat expansion performance datasets

WGS repeat expansion performance was evaluated using data from two sources: individuals from Genomics England (GE) in the 100,000 Genomes Project who were also assessed for expansions using orthogonal approaches, or individuals previously assessed for expansions as part of routine clinical assessment in England. For individuals from the 100,000 Genomes Project, locus-specific PCR tests were obtained from participants that had been screened for REs prior to recruitment to the project, or were identified by WGS as carrying a pathologically expanded allele and subsequently orthogonally confirmed. The GE dataset consisted of 254 patient samples and 634 individual PCR tests. For each individual both alleles at each locus were assessed, with the exception of loci on chromosome X in males, yielding 1,172 normal alleles, 22 premutation alleles and 39 expanded alleles (**Table S2**). Samples utilized by ICSL were drawn from the Genetics Laboratory, at the Cambridge University Hospitals NHS Foundation Trust and included 149 PCR-negative alleles and 160 PCR-positive alleles across 150 patient samples.

**Repeat expansion loci analysed**

Eleven repeats associated with ataxia and late-onset neurodegenerative disorders were tested, including all exonic CAG repeat disorders - Huntington disease (*HTT*; #143100), spinal and bulbar muscular atrophy of Kennedy (*AR*; #313200), dentatorubral-pallidoluysian atrophy (*ATN1*; #125370), spinocerebellar ataxia 1 (*ATXN1*; #164400), spinocerebellar ataxia 2 (*ATXN2*; #183090), Machado-Joseph disease (*ATXN3*; #109150), spinocerebellar ataxia 7 (*ATXN7*; #164500), spinocerebellar ataxia 6 (*CACNA1A;* #183086), and spinocerebellar ataxia 17 (*TBP*; #607136); and two known intronic repeat disorders - frontotemporal dementia and/or amyotrophic lateral sclerosis 1 (*C9orf72*; #105550) and Friedreich ataxia (*FXN*; #229300). Further, we investigated performance for Fragile X syndrome (*FMR1*; #300624) and Myotonic dystrophy 1 (*DMPK*; #*160900*). Genomic coordinates defined for the region of each locus used by ExpansionHunter versions EHv2.5.5 and EHv3.1.2 can be found in **Table S12** and **Table S13** respectively.

**PCR analysis**

Genomic DNA was isolated from peripheral blood leukocytes following standard protocols. PCR-based and Southern blot analysis of repeat expansion loci was performed by the Neurogenetics Laboratory at the National Hospital for Neurology and Neurosurgery (UCLH NHS Trust) for all Genomics England samples. The Genetics Laboratory, Cambridge University Hospitals NHS Foundation Trust assessed all ICSL sequenced samples as part of routine clinical assessment.

**Neurogenetics Laboratory at the National Hospital for Neurology and Neurosurgery**

All PCR assays Southern blot followed accredited diagnostic protocols established laboratory. Allele sizing by fluorescent tethered repeat-primed PCR was previously validated against Sanger-sequenced controls, and control samples of a known repeat expansion size were included in each assessment.

SCA1-7 (*ATXN1, ATXN2, ATXN3, CACNA1A, ATXN7*): Fluorescent tethered repeat-primed PCR (RP-PCR) was performed for each locus using the primers listed in **Table S11**, with PCR conditions modified from Cagnoli et al.[1] Large SCA7 expansion that could not be sized by tethered RPPCR were amplified with flanking primers (**Table S11**) and assessed by electrophoresis on a 2% agarose gel against a 1 kb size standard (Promega).

SCA17 (*TBP*): A fluorescent flanking PCR was performed using the primers listed in **Table S11**, based on Nethsinghe et al.[2]

HD (*HTT*): Two fluorescent PCRs were performed, both based on the method published by Warner et al.[3] The first PCR was a tethered RP-PCR interrogating the CAG repeat size. The second PCR was a flanking PCR also capturing an adjacent highly polymorphic 'CCG' repeat. See **Table S11** for primer sequences.

FRDA (*FXN*): A three-primer fluorescent RP-PCR assay was used following the protocol established by Warner et al,[4] with the 3rd 'non-genomic' primer complementary to the tail of the repeat-binding primer, and a 'long PCR' with flanking primers that was analysed both fluorescently and on 2% agarose gel to approximate the size of the pathogenic expansions.[5] See **Table S11** for primer sequences.

SBMA (*AR*): A fluorescent flanking PCR was performed using published primers[6] listed in **Table S11.**

DRPLA (*ATN1*): A tethered fluorescent RP-PCR was used, using the primers listed in **Table S11** and under standard PCR conditions.

FTD/ALS (*C9orf72*): Two fluorescent RP-PCRs were performed, amplifying opposite ends of the repeat, utilising published primers. RP-PCR1 is derived from Renton et al[7] and RP-PCR2 (a tethered RP-PCR) from DeJesus-Hernandez et al.[8] These were complemented by a flanking PCR, using primers from DeJesus-Hernandez et al.[8] See **Table S11** for sequences. Expansions detected by RP-PCR were confirmed and sized by Southern blotting using a 1 kb single copy probe as previously described[9] but using BsU36I restriction enzyme digests that generate a 6.2 kb band for unexpanded alleles, rather than the EcoRI digest used previously that generates an 8 kb band described in the original protocol.

**Repeat expansion from whole genome sequencing**

STR genotyping from whole genome sequencing was performed using ExpansionHunter.[10,11] In brief, ExpansionHunter (EH) aligns reads to a modified segment of the reference genome that can accommodate STRs of any length. The algorithm employs either an ad hoc[10] or graph-based approach.[11] When the STR is shorter than the WGS read length (e.g. 150 bp) the genotype is identified from the realigned reads. When the STR is longer than the read length, its size is estimated from the number of reads that align to the locus plus those that are entirely composed of repeat sequences (i.e. in-repeat reads). We used two versions of EH, EHv3.1.2 and

EHv2.5.5 for the assessment of the performance of the pipeline (**Table S4**). The overall performance of the pipeline was not affected by EH version (**Table S5** and **Table S15**).

### Visual inspection

Visual inspection of complex WGS variant calls, using tools like Integrated Genome Viewer, IGV,[12] is standard practice in most clinical laboratories. Because repeat expansions can include a significant amount of inserted sequence relative to the reference genome, these common visualizations are not adequate for repeat expansion investigation. To address this gap we used a tool that creates a static visualization of the read pileup against an alternative reference constructed from the putative expanded allele identified by EH (**Figure 2C and 2D**). This tool can be downloaded from https://github.com/Illumina/GraphAlignmentViewer and makes it possible to inspect the evidence used by EH to make a genotype call and identify putative false positives. Additionally, it enables rapid visual inspection of data quality by representing low quality (i.e. <Q20) bases as lower case, facilitating identification of genomic regions or samples that may be impacted by poor data quality. From the visualization, a user is able to identify and interrogate sequencing reads that align to the region allowing for an additional assessment of the ExpansionHunter calls, analogous to how IGV is used to visually confirm SNP calls.

### Interpreting pileup plots

To interpret a pileup plot, first it is important to understand how EH estimates repeat sizes from WGS at a given locus. EH scans the WGS BAM file to identify reads that (1) either fully span the repeat (spanning reads); or (2) include the repeat and the flanking sequence on one side of the repeat (flanking reads); or (3) are fully contained in the repeat ("in-repeat" reads, IRR). It does this by creating a dynamic graph reference genome where the repeat is represented by a loop in the graph and the reads are realigned to this dynamic reference (reference EH & EH graph). If the repeat is shorter than the read length of the sequence data (e.g. an expansion in *ATXN2* or *HTT*), the repeat length, there should be some spanning reads and an exact size is identified. To estimate the length of a repeat longer than the read length (e.g *C9orf72* or *FXN*), IRRs are identified and counted. When the repeat length is close to the read length, the size of the repeat is approximated from the flanking reads that partially overlap the repeat and one of the repeat flanks. If the repeat is longer than the read length, its size is estimated from IRR. In-repeat reads anchored by their mate to the repeat region are used to estimate the size of the repeat up to the fragment length.

The pileup plot is a graphic representation of the sequencing reads that align to the repeat region of interest. Within a pileup graph, the reads supporting each genotype are grouped based on (i) the type of reads: spanning, flanking or IRR; (ii) the repeat length supported by each group. Furthermore, sequencing quality of each base of the read is represented by upper case letters for high quality (>Q20 ; error rate less than 1%) bases or lower case base letters for low quality (<Q20 ; error rate >1%) bases.

In our experiences, genotyping errors may occur if reads are classified incorrectly due low quality data or mosaicism but the sequencing reads represented in the pileup plot can be inspected and interrogated to confirm the repeat length predicted by EH. **Figure S1** shows three examples of different pileup plots that support the genotype predicted by the correspondent EH call:

- Case A: A monoallelic expansion smaller than the read length where 9 spanning reads support the shorter repeat (22 repeats in green box) and two spanning reads support the longer repeat (40 repeats in green box). In addition to the spanning reads there are multiple flanking reads that have up to 39 repeats and provide further evidence of the expansion.
- Case B: A monoallelic expansion larger than the read-length where 14 spanning reads support the shorter allele (2 repeats) and ~26 IRRs provide strong evidence for the expansion.
- Case C: A biallelic expansion larger than the read length where there are no spanning reads for a short allele and ~44 IRRs supporting long alleles. We would expect fewer IRRs if there was only a single expansion - note that there are roughly twice as many IRRs in this example was observed for the monoallelic expansion of case B.

**Repeat sizing**

In order to analyze the correlation between PCR and WGS repeat-size estimates, genomes for which PCR exact lengths were available were taken into account. A total of 418 PCR tests were analyzed corresponding to 805 alleles (marked as `Yes` in `repeat_sizing_test` column in **Table S4**). In this analysis, we assumed that the PCR call is correct. **Table S4** provides the allele sizes from PCR alone and the repeat-size estimates of both EH versions. The overall concordance and discordance computation and by locus is shown in **Table S6**. The EH repeat lengths were in agreement with the original PCR estimates for 92.5% of the alleles, taking into account 1 PCR repeat error across all loci (**Figure2B**; **Figure S4**).

**100,000 Genomes Project**

Following ethical approval (14/EE/1112), consenting participants with unmet diagnostic need in rare diseases after usual care testing (single gene tests, arrays or exomes) were recruited in partnership with hospitals across England. Probands and, where feasible, other family members were enrolled by multiple specialties according to eligibility criteria (see below). Standardised baseline clinical data were recorded using the Human Phenotyping Ontology, HPO,[13] and whole blood was drawn for DNA extraction followed by whole genome sequencing. Families were analysed using the virtual gene panel relevant to the disease of their recruitment. In addition, reflecting the complexity of many clinical presentations, additional gene panels were used based on the HPO terms. For example, a child recruited under the 'Intellectual disability' category who also has dystonia, would be analysed with both the 'Intellectual disability' and 'Early onset dystonia ' gene panels.

**Inclusion criteria for testing 100,000 Genomes Project participants using the WGS pipeline to detect repeat expansions**

Four subgroups of participants with a neurological phenotype potentially consistent with a repeat expansion disorder caused by one of the following loci *AR*, *ATN1*, *ATXN1*, *ATXN2*, *ATXN3*, *CACNA1A*, *ATXN7*, *C9orf72*, *DMPK*, *FXN*, *FMR1*, *HTT* and *TBP* were selected as follows: Subgroup A included individuals recruited under any of the following diseases: amyotrophic lateral sclerosis or motor neuron disease, Charcot-Marie-Tooth disease, early onset dementia, early onset dystonia, complex parkinsonism, hereditary ataxia, hereditary spastic paraplegi,early onset and familial Parkinson's disease. For all these diseases only adults (i.e. individuals equal or older than 18 years old) were selected except for hereditary ataxia, where children were also included. Furthermore, participants recruited under ultra-rare undescribed monogenic disorders (i.e. participants that did not fit clinically a specific eligible disease) and whose HPO terms were suggestive of ataxia or a neurodegenerative disorder were also included. Subgroup A participants were selected if they were analysed using any of the following panels: amyotrophic lateral sclerosis/motor neuron disease, hereditary neuropathy, early onset dementia (encompassing fronto-temporal dementia and prion disease), Parkinson disease and complex parkinsonism, early onset dystonia, hereditary spastic paraplegia or hereditary ataxia. Expansions in *AR, ATN1, ATXN1, ATXN2, ATXN3, ATXN7, CACNA1A, C9orf72, FXN, HTT,* or *TBP* were analysed using the pathogenic expansion cutoffs defined in **Table S4,** with the exception of *HTT* where the the premutation cutoff was used.

Subgroup B included individuals that were assigned the intellectual disability disease panel and genetic epilepsy syndromes, congenital muscular dystrophy, hereditary ataxia, hereditary spastic paraplegia, mitochondrial disorders, inherited white matter disorders, optic neuropathy, or brain channelopathy. In this group repeat expansions in *ATN1, ATXN1, ATXN2, ATXN3, ATXN7, HTT*, and *TBP* using the full mutation cutoffs defined in **Table S4** were investigated.

Subgroup C included participants assigned to any of the following diseases: intellectual disability, Kabuki syndrome, congenital muscular dystrophy, congenital myopathy, skeletal muscle channelopathies, distal myopathies. Genomes for individuals included in this dataset were assessed for expansions in *DMPK* using the cutoffs described in **Table S4**.

Finally, Subgroup D included children (i.e. individuals younger than 18 years old in 2020) having been assigned Intellectual disability or Kabuki syndrome disease panels. These individuals were assessed for expansions in *FMR1* using the premutation threshold defined in **Table S4**.

For all these four subgroups the total number of RE detected before and after visual inspection is presented in **Table S8**.

**100,000 Genomes Project eligibility statements**

Participants were recruited to the 100,000 Genomes Project according to the eligibility criteria for conditions approved within the Genomics England Rare Diseases Programme. Eligibility criteria for the conditions analysed in this study are listed below. For further information, please refer to "Rare Disease Conditions Eligibility Criteria - 100,000 Genomes Project".[14]

**CHARCOT-MARIE-TOOTH DISEASE**

**Inclusion criteria**

- Unexplained peripheral neuropathy affecting motor, sensory or autonomic nerves progressing over >2 years +/- additional neurological signs.

**Exclusion criteria**

- History of trauma
- Known acquired metabolic, vascular, inflammatory or immunological cause - History of alcohol excess
- Evidence of malignancy
- ENG/EMG suggest acquired pathology

**Prior genetic testing guidance**

- Results should have been reviewed for all genetic tests undertaken, including disease-relevant genes in exome sequencing data. The patient is not eligible if they have a molecular diagnosis for their condition.
- Genetic testing should continue according to routine local practice for this phenotype regardless of recruitment to the project; results of these tests must be submitted via the 'Genetic investigations' section of the data capture tool to allow comparison of WGS with current standard testing.
- PLEASE NOTE: The sensitivity of WGS compared to current diagnostic genetic testing has not yet been established. It is therefore important that tests which are clinically indicated under local standard practice continue to be carried out.

**Prior genetic testing genes**

- Testing for the chromosome 17p11.2 duplication is strongly recommended PRIOR TO RECRUITMENT as this may not be reliably detected by WGS using current analysis techniques; other tests below should be considered where this is in line with current local practice including:
- *PMP22* point mutations, *GJB1, MPZ, MFN2* (*MFN2* axonal only)

**EARLY ONSET DEMENTIA**

**Early onset dementia inclusion criteria**

- Progressive cognitive deterioration with change in memory, vision, behaviour or language with functional impairment
- Age at onset <60 years OR
- Later onset with family history of dementia of the same type in a first or second degree relative
- Individuals with severe or syndromic disease should be recruited according to standard guidance, typically as trios. Disease status of apparently unaffected participants should be determined according to standard clinical practice to detect cryptic disease. In other cases, unaffected individuals should not be recruited. Recruitment in such families should favour multiplex families over single isolated cases. These singleton recruits will not contribute to the overall singleton monitoring metrics applied to GMCs.

**Early onset dementia exclusion criteria**

- Identified underlying cause, e.g. structural brain lesion. NB in uncertain cases with anxiety/depression brain atrophy on imaging, CSF findings or EEG abnormalities should be available to support the diagnosis of a primary degenerative syndrome

**Prior genetic testing guidance**

- Results should have been reviewed for all genetic tests undertaken, including disease-relevant genes in exome sequencing data. The patient is not eligible if they have a molecular diagnosis for their condition.
- Genetic testing should continue according to routine local practice for this phenotype regardless of recruitment to the project; results of these tests must be submitted via the 'Genetic investigations' section of the data capture tool to allow comparison of WGS with current standard testing.
- PLEASE NOTE: The sensitivity of WGS compared to current diagnostic genetic testing has not yet been established. It is therefore important that tests which are clinically indicated under local standard practice continue to be carried out.

**Early onset Dementia prior genetic testing genes**

- Testing of the following genes should be carried out PRIOR TO RECRUITMENT where this is in line with current local practice:
    - Clinical syndrome Alzheimer disease: *PSEN1, APP*
    - Clinical syndrome FTLD: *MAPT, C9ORF72, GRN*
    - Clinical syndrome Prion disease: *PRNP*

- PLEASE NOTE: The sensitivity of WGS compared to current diagnostic genetic testing has not yet been established. It is therefore important that tests which are clinically indicated under local standard practice continue to be carried out.

**EARLY ONSET DYSTONIA**

**Early onset dystonia inclusion criteria**

- Dystonia affecting any body part, usually spreading to involve multiple body regions (e.g. multifocal, segmental, generalised)
- Age at onset <31 years or later onset with family history of early onset dystonia
- May be paroxysmal/episodic dystonia
- May be associated with myoclonus as in myoclonic dystonia
- This disease category includes dopa responsive dystonia.

**Early onset dystonia exclusion criteria**

- Underlying cause for clinical syndrome identified, e.g. cerebral palsy, structural brain lesion, Wilson disease, psychogenic dystonia

**Prior genetic testing guidance**

- Results should have been reviewed for all genetic tests undertaken, including disease-relevant genes in exome sequencing data. The patient is not eligible if they have a molecular diagnosis for their condition.
- Genetic testing should continue according to routine local practice for this phenotype regardless of recruitment to the project; results of these tests must be submitted via the

‘Genetic investigations’ section of the data capture tool to allow comparison of WGS with current standard testing.

- PLEASE NOTE: The sensitivity of WGS compared to current diagnostic genetic testing has not yet been established. It is therefore important that tests which are clinically indicated under local standard practice continue to be carried out.

**Early onset dystonia prior genetic testing genes**

- Testing of the following genes should be carried out PRIOR TO RECRUITMENT where this is in line with current local practice:
    - *TOR1A*

**COMPLEX PARKINSONISM (INCLUDES PALLIDO-PYRAMIDAL SYNDROMES)**

**Complex Parkinsonism inclusion criteria**

- Progressive motor syndrome with parkinsonism (bradykinesia with one of tremor, gait disorder, stiffness)
- Additional features may include spasticity, gaze palsy, early dementia, early bulbar failure, dyspraxia, ataxia, postural hypotension, cortical sensory loss, brain iron accumulation on MRI brain
- Aat onset <= 45 years or later onset with family history of similar condition in other family members
- Individuals with severe or syndromic disease should be recruited according to standard guidance, typically as trios. Disease status of apparently unaffected participants should be determined according to standard clinical practice to detect cryptic disease. In other

cases, unaffected individuals should not be recruited. Recruitment in such families should favour multiplex families over single isolated cases. These singleton recruits will not contribute to the overall singleton monitoring metrics applied to GMCs.

**Complex Parkinsonism exclusion criteria**

- Underlying cause not identified, e.g. structural brain lesion, Wilson disease

**Prior genetic testing guidance**

- Results should have been reviewed for all genetic tests undertaken, including disease-relevant genes in exome sequencing data. The patient is not eligible if they have a molecular diagnosis for their condition.
- Genetic testing should continue according to routine local practice for this phenotype regardless of recruitment to the project; results of these tests must be submitted via the 'Genetic investigations' section of the data capture tool to allow comparison of WGS with current standard testing.
- PLEASE NOTE: The sensitivity of WGS compared to current diagnostic genetic testing has not yet been established. It is therefore important that tests which are clinically indicated under local standard practice continue to be carried out.

**Complex Parkinsonism prior genetic testing genes**

- Testing of the following genes should be carried out PRIOR TO RECRUITMENT where this is in line with current local practice:
- *C9ORF72, GRN, MAPT* in cases with a clinical presentation suggestive of cortico-basal/PSP syndrome

### HEREDITARY ATAXIA

#### Hereditary ataxia inclusion criteria

- Unexplained cerebellar ataxia progressing over >2 years +/- spasticity, peripheral neuropathy, or bulbar dysfunction.
- Individuals with syndromic disease or disease onset <30 years should be recruited according to standard guidance, typically as trios. Disease status of apparently unaffected participants should be determined according to standard clinical practice to detect cryptic disease.
- In other cases, unaffected individuals should not be recruited. Recruitment in such families should favour multiplex families over single isolated cases. These singleton recruits will not contribute to the overall singleton monitoring metrics applied to GMCs.

#### Hereditary ataxia exclusion criteria

- No structural or inflammatory (MS-like) lesions on brain MRI. - No history of alcohol excess.
- Normal thyroid function.
- No evidence of malignancy.

#### Prior genetic testing guidance

- Results should have been reviewed for all genetic tests undertaken, including disease-relevant genes in exome sequencing data. The patient is not eligible if they have a molecular diagnosis for their condition.
- Genetic testing should continue according to routine local practice for this phenotype regardless of recruitment to the project; results of these tests must be submitted via the

‘Genetic investigations’ section of the data capture tool to allow comparison of WGS with current standard testing.

- PLEASE NOTE: The sensitivity of WGS compared to current diagnostic genetic testing has not yet been established. It is therefore important that tests which are clinically indicated under local standard practice continue to be carried out.

**Hereditary ataxia prior genetic testing genes**

- Testing for genes which are affected by trinucleotide repeats is strongly recommended PRIOR TO RECRUITMENT as these will not be reliably detected by WGS using current analysis techniques including: common trinucleotide repeat disorders excluded (*ATXN1, ATXN2, ATXN3, CACNA1A, ATXN7, TBP, ATN1, FXN* (only recessive history), *FMR1*)

**HEREDITARY SPASTIC PARAPLEGIA**

**Hereditary spastic paraplegia inclusion criteria**

- Unexplained spastic paraplegia progressing over >2 years +/-, peripheral neuropathy, or ataxia.
- Individuals with syndromic disease or disease onset <30 years should be recruited according to standard guidance, typically as trios. Disease status of apparently unaffected participants should be determined according to standard clinical practice to detect cryptic disease.
- In other cases, unaffected individuals should not be recruited. Recruitment in such families should favour multiplex families over single isolated cases. These singleton recruits will not contribute to the overall singleton monitoring metrics applied to GMCs.

**Hereditary spastic paraplegia exclusion criteria**

- No structural or inflammatory (MS-like) lesions on brain MRI.

**Prior genetic testing guidance**

- Results should have been reviewed for all genetic tests undertaken, including disease-relevant genes in exome sequencing data. The patient is not eligible if they have a molecular diagnosis for their condition.
- Genetic testing should continue according to routine local practice for this phenotype regardless of recruitment to the project; results of these tests must be submitted via the 'Genetic investigations' section of the data capture tool to allow comparison of WGS with current standard testing.
- PLEASE NOTE: The sensitivity of WGS compared to current diagnostic genetic testing has not yet been established. It is therefore important that tests which are clinically indicated under local standard practice continue to be carried out.

**Hereditary spastic paraplegia prior genetic testing genes**

- Testing of the following genes should be carried out PRIOR TO RECRUITMENT where this is in line with current local practice:
    - *SPAST, ATL1*
    - Normal very long chain fatty acid studies

**EARLY ONSET AND FAMILIAL PARKINSON'S DISEASE**

**Early onset and familial Parkinson's Disease inclusion criteria**

- Early onset (<= 45 years of age) or history of other family member with Parkinson’s Disease
- Bradykinesia plus at least one of rigidity, rest tremor and gait disturbance - May have concurrent dystonia (common in early onset PD)
- May have positive family history or consanguinity
- If complex features, e.g. spasticity, early dementia, gaze palsy, Neurodegeneration with Brain Iron Accumulation, please recruit to Complex Parkinsonism
- May develop Lewy Body/PD type dementia
- Individuals with severe or syndromic disease should be recruited according to standard guidance, typically as trios. Disease status of apparently unaffected participants should be determined according to standard clinical practice to detect cryptic disease. In other cases, unaffected individuals should not be recruited. Recruitment in such families should favour multiplex families over single isolated cases. These singleton recruits will not contribute to the overall singleton monitoring metrics applied to GMCs.

**Early onset and familial Parkinson's Disease exclusion criteria**

- Underlying cause for clinical syndrome identified, e.g. cerebral palsy, dopa-responsive dystonia, structural brain lesion, Wilson disease, psychogenic dystonia

**Prior genetic testing guidance**

- Results should have been reviewed for all genetic tests undertaken, including disease-relevant genes in exome sequencing data. The patient is not eligible if they have a molecular diagnosis for their condition.

- Genetic testing should continue according to routine local practice for this phenotype regardless of recruitment to the project; results of these tests must be submitted via the 'Genetic investigations' section of the data capture tool to allow comparison of WGS with current standard testing.
- PLEASE NOTE: The sensitivity of WGS compared to current diagnostic genetic testing has not yet been established. It is therefore important that tests which are clinically indicated under local standard practice continue to be carried out.

**AMYOTROPHIC LATERAL SCLEROSIS OR MOTOR NEURON DISEASE**

**Amyotrophic lateral sclerosis or motor neuron disease inclusion criteria**

- Progressive upper and/or lower motor neuron disease degeneration with clinical features of amyotrophy, spasticity, bulbar/pseudo-bulbar involvement
- EMG/NCS consistent with MND
- Positive family history of other affected family members with ALS or with FTD/ALS like phenotype or disease onset below 40 years.
- Individuals with severe or syndromic disease should be recruited according to standard guidance, typically as trios. Disease status of apparently unaffected participants should be determined according to standard clinical practice to detect cryptic disease. In other cases, unaffected individuals should not be recruited. Recruitment in such families should favour multiplex families over single isolated cases. These singleton recruits will not contribute to the overall singleton monitoring metrics applied to GMCs.

**Amyotrophic lateral sclerosis or motor neuron disease exclusion criteria**

- Identified underlying cause for clinical syndrome e.g. multi-focal motor neuropathy, lymphoma

**Prior genetic testing guidance**

- Results should have been reviewed for all genetic tests undertaken, including disease-relevant genes in exome sequencing data. The patient is not eligible if they have a molecular diagnosis for their condition.
- Genetic testing should continue according to routine local practice for this phenotype regardless of recruitment to the project; results of these tests must be submitted via the 'Genetic investigations' section of the data capture tool to allow comparison of WGS with current standard testing.
- PLEASE NOTE: The sensitivity of WGS compared to current diagnostic genetic testing has not yet been established. It is therefore important that tests which are clinically indicated under local standard practice continue to be carried out.

**Amyotrophic lateral sclerosis or motor neuron disease prior genetic testing genes**

- Testing of the following genes should be carried out PRIOR TO RECRUITMENT where this is in line with current local practice: *C9ORF72, SOD1*

**INTELLECTUAL DISABILITY**

**Intellectual disability inclusion criteria**

- Moderate to Severe/ Profound ID disproportionate to parental IQ unless the family history is consistent with an X- linked disorder
- Congenital onset

- Developmental Delay
- +/- clinical features suggestive of a specific syndrome - Metabolic causes have been excluded

**Intellectual disability exclusion criteria**

- Antenatal history suggestive of non-genetic cause
- Proven congenital or neonatal infections
- Known genetic cause already identified
- Microarray analysis abnormal and clearly pathogenic

**Prior genetic testing guidance**

- Results should have been reviewed for all genetic tests undertaken, including disease-relevant genes in exome sequencing data. The patient is not eligible if they have a molecular diagnosis for their condition.
- Genetic testing should continue according to routine local practice for this phenotype regardless of recruitment to the project; results of these tests must be submitted via the 'Genetic investigations' section of the data capture tool to allow comparison of WGS with current standard testing.

**Intellectual disability prior genetic testing genes**

- Testing of the following genes should be carried out PRIOR TO RECRUITMENT where this is in line with current local practice:
- For syndromes where the cause of disease is 1-2 genes these need to be excluded before Genomics England recruitment, e.g. for Kabuki syndrome, *MLL2* (*KMT2D*), and *KDM6A* should have been tested

### CONGENITAL MUSCULAR DYSTROPHY

#### Congenital muscular dystrophy inclusion criteria

- Muscle weakness with onset in infancy or early childhood AND
- elevated creatine kinases or muscle biopsy with dystrophic changes - Availability of CK and muscle biopsy results
- Dystrophic changes on muscle biops
- Congenital muscular dystrophy exclusion criteria

#### Prior genetic testing guidance

- Results should have been reviewed for all genetic tests undertaken, including disease-relevant genes in exome sequencing data. The patient is not eligible if they have a molecular diagnosis for their condition.
- Genetic testing should continue according to routine local practice for this phenotype regardless of recruitment to the project; results of these tests must be submitted via the ‘Genetic investigations’ section of the data capture tool to allow comparison of WGS with current standard testing.

### CONGENITAL MYOPATHY

Relevant diseases:

- Congenital myopathy

#### Congenital myopathy inclusion criteria

- Muscle weakness
- one or more of the following histopathological features
- type 1 predominance or uniformity - congenital fibre type disproportion - central cores
- Multi-minicores
- nemaline rods - central nuclei
- Availability of CK, muscle CT/MR imaging, muscle biopsy and neurophysiological studies

**Congenital myopathy exclusion criteria**

- Absence of muscle weakness
- CK more than 5x normal
- dystrophic features on muscle biopsy

**Prior genetic testing guidance**

- Results should have been reviewed for all genetic tests undertaken, including disease-relevant genes in exome sequencing data. The patient is not eligible if they have a molecular diagnosis for their condition.
- Genetic testing should continue according to routine local practice for this phenotype regardless of recruitment to the project; results of these tests must be submitted via the 'Genetic investigations' section of the data capture tool to allow comparison of WGS with current standard testing

**DISTAL MYOPATHIES**

**Distal myopathies inclusion criteria**

- Unexplained predominantly distal muscle weakness, onset at any age - Acquired myopathies excluded by relevant clinical investigations
- Serum creatine kinase (CK) assessment
- Muscle Biopsy with immunohistochemistry (IH)
- Neurophysiology performed
- Muscle MRI (optional)
- Dried blood spot test for Pompe disease performed

**Distal myopathies exclusion criteria**

NA

**Prior genetic testing guidance**

- Results should have been reviewed for all genetic tests undertaken, including disease-relevant genes in exome sequencing data. The patient is not eligible if they have a molecular diagnosis for their condition.
- Genetic testing should continue according to routine local practice for this phenotype regardless of recruitment to the project; results of these tests must be submitted via the 'Genetic investigations' section of the data capture tool to allow comparison of WGS with current standard testing.

**Distal myopathies prior genetic testing genes**

Testing of the following genes should be carried out PRIOR TO RECRUITMENT where this is in line with current local practice:

- *DMD* analysis by MLPA or equivalent
- *DUX1* and *DMPK* exclusion by conventional genetic testing

- Exclusion by genetic testing of any gene indicated by IH.
- In the presence of evidence of myofibrillar myopathy on muscle biopsy IH exclusion of *LDB3, MYOT, DES, CRYAB* by sequencing

**SKELETAL MUSCLE CHANNELOPATHIES**

**Skeletal Muscle Channelopathies inclusion criteria**

- Episodic flaccid paralysis or weakness and/or myotonia
- May develop progressive, usually proximal, weakness
- Electrophysiology including long and short exercise testing - Intra-attack potassium documented whenever possible
- Normal renal function and thyroid function

**Skeletal Muscle Channelopathies exclusion criteria**

- Primary renal or endocrine problem that may be causative - Associated loss of consciousness with attacks

**Prior genetic testing guidance**

- Results should have been reviewed for all genetic tests undertaken, including disease-relevant genes in exome sequencing data. The patient is not eligible if they have a molecular diagnosis for their condition.
- Genetic testing should continue according to routine local practice for this phenotype regardless of recruitment to the project; results of these tests must be submitted via the 'Genetic investigations' section of the data capture tool to allow comparison of WGS with current standard testing.

**Skeletal Muscle Channelopathies prior genetic testing genes**

Testing of the following genes should be carried out PRIOR TO RECRUITMENT where this is in line with current local practice:

- Myotonia: *DMPK, CNBP, SCN4A, CLCN1* (including MLPA) - Episodic weakness: *CACNA1S, SCN4A, KCNJ2*
